# M-CSF induces a coordinated myeloid and NK cell differentiation program protecting against CMV after hematopoietic cell transplantation

**DOI:** 10.1101/2023.01.30.526105

**Authors:** Prashanth K. Kandalla, Julien Subburayalu, Clément Cocita, Bérengère de Laval, Elena Tomasello, Johanna Iacono, Jessica Nitsche, Maria M. Canali, Wilfried Cathou, Gilles Bessou, Noushine Mossadegh-Keller, Caroline Huber, Sandrine Sarrazin, Guy Mouchiroud, Roland Bourette, Marie-France Grasset, Marc Dalod, Michael H. Sieweke

## Abstract

Immunosuppressed patients are highly susceptible to viral infections. Therapies reconstituting autologous antiviral immunocompetence could therefore represent an important prophylaxis and treatment. Herpesviridae including cytomegalovirus (CMV) are a major cause of morbidity and mortality in patients after hematopoietic cell transplantation (HCT). Here, we show in a mouse model of HCT that macrophage colony-stimulating factor (M-CSF/CSF-1), a key cytokine for myeloid and monocytic differentiation, promoted rapid antiviral activity and protection from viremia caused by murine CMV. Mechanistically, M-CSF stimulated a coordinated myeloid and natural killer (NK) cell differentiation program culminating in increased NK cell numbers and production of granzyme B and interferon-γ. This NK cell response depended upon M-CSF-induced myelopoiesis leading to IL15Rα-mediated presentation of IL-15 on monocytes. Furthermore, M-CSF also induced differentiation of plasmacytoid dendritic cells producing type I interferons, which supported IL-15-mediated protection. In the context of human HCT, M-CSF induced monopoiesis, increased IL15Rα expression on monocytes and elevated numbers of functionally competent NK cells in G-CSF-mobilized human hematopoietic stem and progenitor cells. Together, our data show that M-CSF induces an integrated multistep differentiation program that culminates in increased NK cell numbers and activation, thereby protecting graft recipients from CMV infection. Thus, our results identify a mechanism by which M-CSF-induced myelopoiesis can rapidly reconstitute antiviral activity during leukopenia following HCT.

**Key points:**

- M-CSF protects from lethal CMV viremia during leukopenia following hematopoietic cell transplantation, a vulnerable period of immunosuppression.
- Early action of M-CSF on donor hematopoietic stem and progenitor cells rapidly reconstitutes antiviral immune responses.
- M-CSF stimulates a coordinated myeloid-NK cell-differentiation program resulting in increased NK cell numbers and activity.
- Increased NK cell differentiation and activity depends on M-CSF-induced myelopoiesis generating IL-15-producing monocytes and I-IFN-producing pDCs.
- M-CSF also stimulates monopoiesis, IL15Ra expression in monocytes and functional NK cell differentiation in G-CSF-mobilized human PBMC.
- No impaired HCT engraftment or proclivity to graft-versus-host-disease by M-CSF.
- M-CSF could provide a single cytokine therapy addressing a major medical need, supporting current antiviral therapies during leukopenia following HCT.

**Visual abstract:** 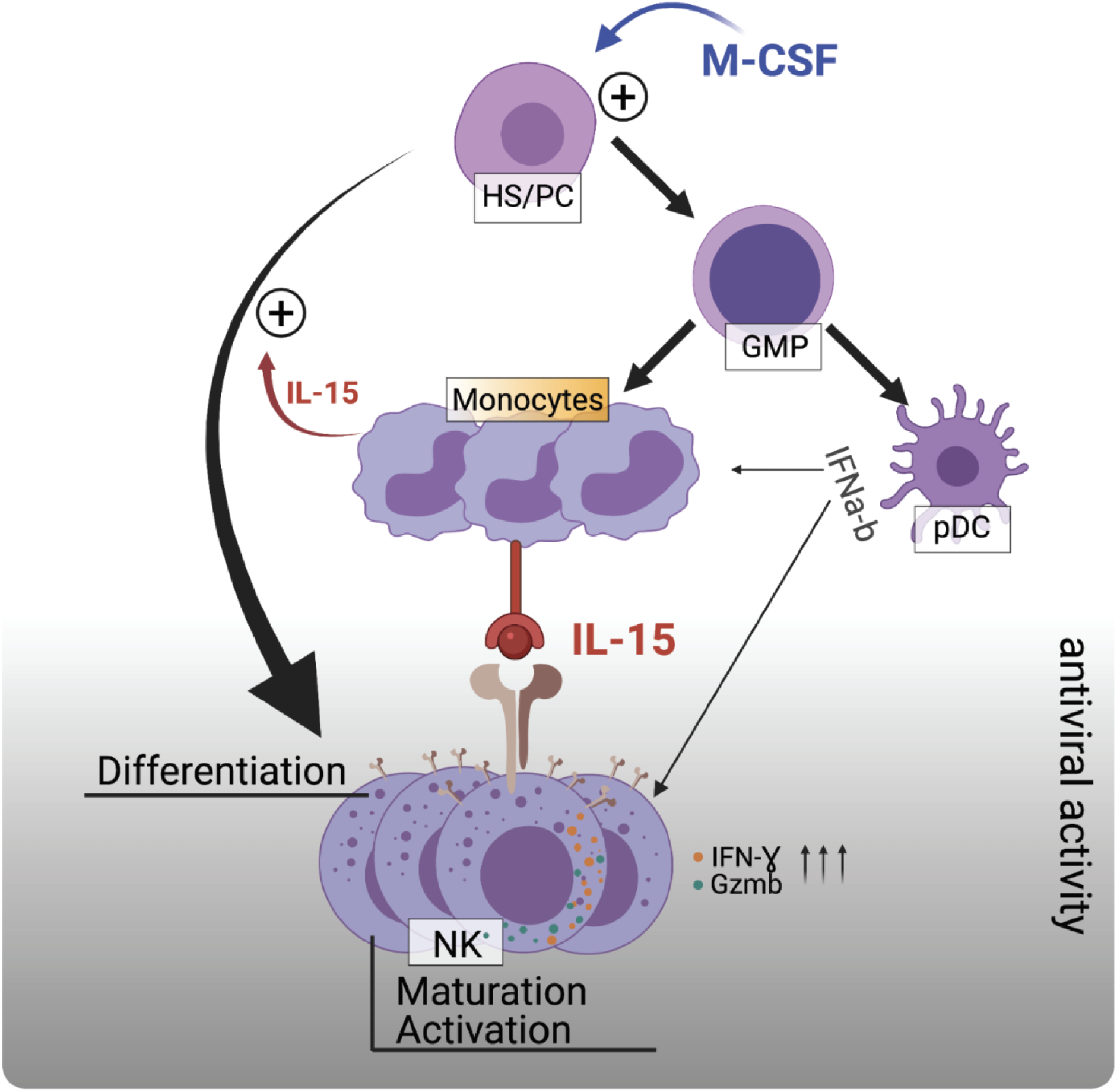

**One Sentence Summary:** M-CSF drives myeloid reconstitution to support CMV-directed natural killer cell competence via IL-15/I-IFN after hematopoietic cell transplantation.

## INTRODUCTION

The first months after hematopoietic cell transplantation (HCT) are characterized by profound immunosuppression, which leaves patients at high risk of viral infection or reactivation of common opportunistic viruses such as cytomegalovirus (CMV). The infection itself but also its subsequent treatment is associated with significant morbidity and mortality (*1–5*). Although vaccines against CMV are under development, they are not yet routinely available in the clinic (*6*). Moreover, antiviral treatments based on inhibition of viral replication are limited to specific viruses, can have significant bone marrow toxicity, and run the risk of variant development and breakthrough infections (*2–5, 7*). Cell-based therapies are still not widely on-hand and associated with high costs (*8*). Biologics stimulating the patient’s general antiviral immune response could therefore be a welcome alternative or complementary treatment option but are currently unavailable.

Myeloid cytokines can massively alter hematopoietic output (*9*) but G-CSF, the major factor in clinical use, has no effect on antiviral immunity (*10*). This appears likely because G-CSF confers its activity only on late myeloid progenitors and mature myeloid cells. By contrast, M-CSF, another myeloid cytokine released during infections (*11–13*), and known to promote myelopoiesis (*14–16*), can directly act on hematopoietic stem and progenitor cells (HSPCs) to induce emergency myelopoiesis (*13*). Importantly, in concert with the myeloid transcription factor MafB, M-CSF selectively controls asymmetric myeloid commitment division in HSPCs (*17, 18*). Consequently, M-CSF stimulates myeloid cell production without exhausting HSPCs (*13, 19*). M-CSF can protect against bacterial and fungal infections after HCT (*19*). However, antiviral activities of M-CSF have not been reported yet.

CMV can lead to a diverse range of pathologies in immunocompromised humans (*20, 21*) and the closely related murine CMV (MCMV) has similar cellular tropism and kinetics (*22, 23*). The spleen is an early site for filtering blood-borne virus and initiated immune responses, whereas the liver is a principal site of viral infection after its decline in the spleen (*24*). Type I interferons (I-IFNs) (*25*), produced by plasmacytoid dendritic cells (pDCs) (*26, 27*), constitute a first line of defense against CMV with natural killer (NK) cells and cytotoxic T cells coming in as a critical second and third wave of the immune response that block viral replication by killing infected cells (*28*). Cytokines including IL-12 and IL-15 produced by conventional dendritic cells (cDCs) can indirectly contribute to viral defense by stimulating NK cell proliferation, activation, and effector function (*25, 29–31*). Other myeloid cells have been shown to have indirect and diverse roles in the response to CMV infection. Whereas Ly6C^-^CX3CR1^+^ patrolling monocytes act as carriers of CMV and can disseminate viral infection to distant organs throughout the body (*32*), Ly6C^+^CCR2^+^ inflammatory monocytes can activate NK and cytotoxic memory CD8^+^ T cells during microbial infection, including MCMV (*33, 34*). Culture models proved that the ability of macrophages to resist MCMV infection depends on signaling mechanisms via I-IFNs and type II IFNs (II-IFNs) (*35–37*), which might also be important *in vivo*. Myeloid-specific deletion of signal transducer and activator of transcription (STAT)1, a key transcription factor for mounting IFN responses, is also required for the early control of MCMV infection and spleen pathology but does not affect viral clearance (*38*). Hence, the role of myeloid cells in MCMV infection appears multifaceted and complex.

Interestingly, in the myeloid STAT1 deletion model, the ability to combat early MCMV infection correlated with the ability to mount extramedullary hematopoiesis (*38*). In this study we have specifically investigated the role of emergency hematopoiesis on MCMV infection under leukopenic conditions and report that M-CSF-induced myelopoiesis promotes rapid reconstitution of antiviral activity and protection from infection. Using a murine model of HCT and infection with lethal doses of MCMV, we observed that M-CSF treatment prompted antiviral immunity resulting in substantially improved survival and pathogen clearance in mice.

Dissecting the mechanism underlying this M-CSF-mediated protection against MCMV infection, we identified a multistep differentiation program in which M-CSF-induced myelopoiesis further stimulated NK cell differentiation and activation via IL-15 and I-IFN mediators. Lastly, we observed that M-CSF also induced intermediate monocyte differentiation from human G-CSF-mobilized HSPCs, enhanced IL15Rα expression on monocytes and increased functional NK cells numbers.

## RESULTS

### M-CSF protects HCT recipients from CMV viremia and mortality

CMV infection/reactivation remains a perilous threat during immunosuppression (*1, 39, 40*). MCMV is a natural pathogen in mice that recapitulates pathomechanisms of human CMV infection (*23*). To study the antiviral effects of M-CSF on MCMV under leukopenic conditions, we used a murine HCT model (*19*). As shown in Fig. 1A, mice received three injections of murine M-CSF (*41*) or PBS at the time of HCT and were infected 14 days later with MCMV doses accounting for 80-90% lethality in untreated transplant recipients (fig. S1A). Survival rates significantly increased, from 25% to 81.8% in M-CSF-treated mice (Fig. 1B). Mice receiving four treatments of murine M-CSF over several days (fig. S1B) or human M-CSF (fig. S1C) both showed improved survival rates. Accordingly, we used three treatments at the time of transplant throughout the study, although singular M-CSF-treatment improved survival rates (fig. S1D). M-CSF-treated mice showed less severe liver injury with a proclivity for scarcer inflammatory foci (Fig. 1C), a reduction of apoptotic or necrotic hepatocytes (Fig. 1D) and decreased necrotic areas after MCMV infection (Fig. 1E). M-CSF-treated mice also showed a decreased viral load as shown by reduced number of infected hepatocytes (Fig. 1F), viral protein IE1 (Fig. 1G) and viral RNA copy numbers (Fig. 1H).

**Fig. 1.**
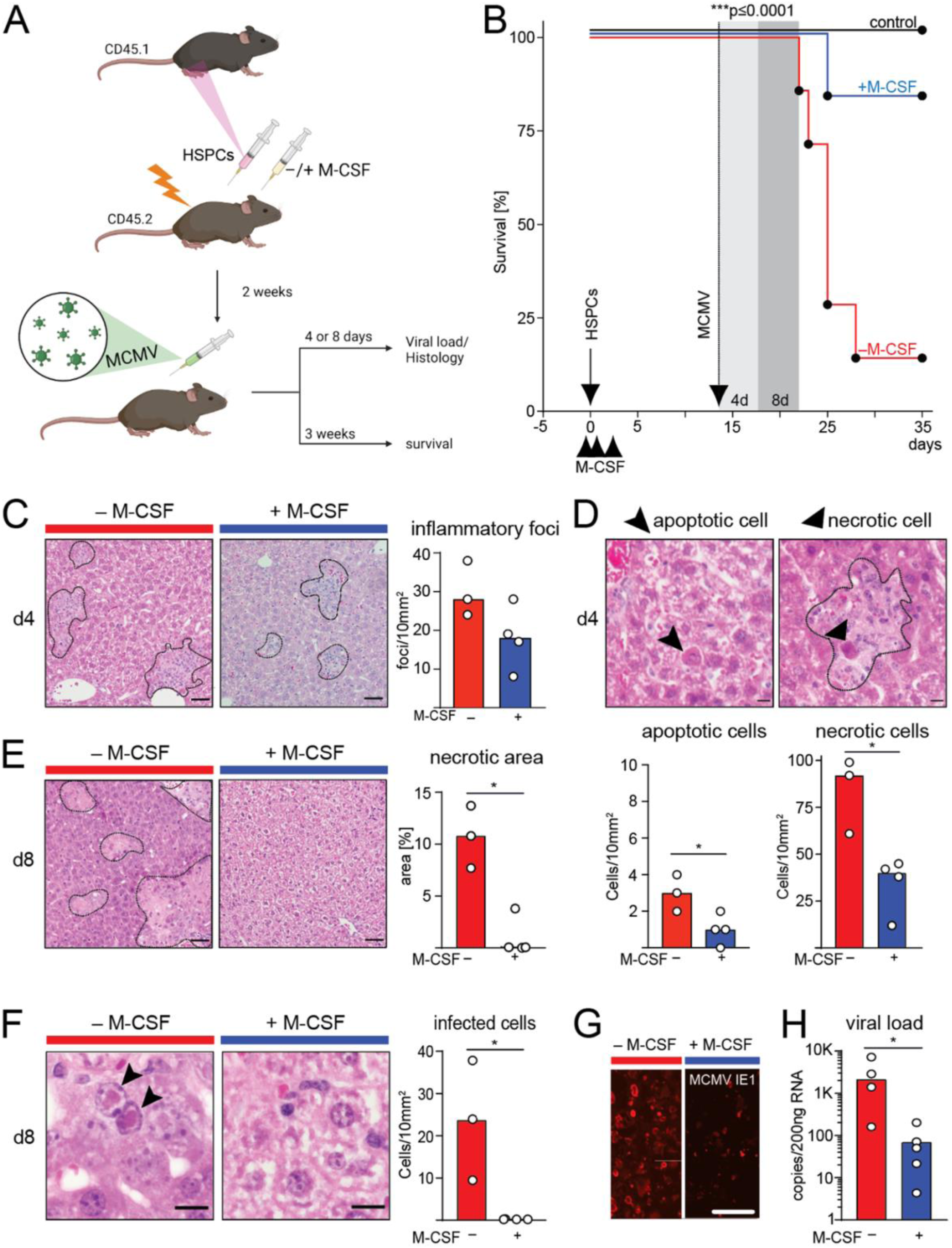
M-CSF protects HSPC recipients from CMV viremia and mortality. (A) Leukopenia model to study MCMV viremia. (B) Survival of mice after HSPC transplantation (arrow), MCMV infection (stippled arrow) and treatment (arrowheads) with control PBS (-M-CSF; n = 12) or three doses of 10µg mouse recombinant M-CSF (+M-CSF; n = 11). Transplanted, uninfected mice (n = 5) are shown as control. (C) Histopathology of MCMV-induced hepatitis. Assessment of inflammatory foci 4 days after infection of transplanted mice treated with M-CSF or control PBS. Example of hematoxylin and eosin (H&E)-staining and inflammatory foci (n = 4). (D) Histopathology of MCMV-induced hepatitis. Apoptotic (arrowheads) and necrotic (arrows) hepatocytes and quantification as median cell numbers per area (n = 4). (E) Histopathology of MCMV-induced hepatitis. Assessment of necrotic area 8 days after infection of transplanted mice (H&E). Percentage of affected areas (n = 4). (F) Histological analysis of infected hepatocytes (H&E); quantification per area (n = 4). (G) Immunofluorescence staining of MCMV E1 protein. (H) RT-qPCR-based quantitation of viral mRNA per 200 ng RNA (n = 5). \*\*\**P* < 0.0001 by Mantel-Cox test (B). \**P* < 0.05 by two-tailed Mann-Whitney *U*-test (C-G). All data are representative of at least two independent experiments.

Together, these results demonstrated that M-CSF treatment protected HCT recipients from MCMV-induced tissue damage and lethality.

### M-CSF treatment increases NK cell abundance, differentiation, and activation

Since NK cells are early antiviral effector cells, including during HCT (*42*), we investigated whether M-CSF treatment influenced NK cells. We observed an increase in NK cell numbers in the spleen two weeks after M-CSF treatment both in uninfected mice and after infection (Fig. 2A). Separate analysis of CD45.2^+^ recipient and CD45.1^+^ graft donor cells revealed that most of the NK cell increase arose from donor cells (fig. S2A). Since M-CSF is short-lived (*43*), but increased NK cell numbers two weeks after application, we posed the mechanism to act on NK cell progenitors. NK cell differentiation stages can be identified by differential expression of surface markers and transcription factors (Fig. 2B) (*44–46*). NK cell progenitors express CD122, CD27 and NKG2D but not the mature markers NK1.1 and NKp46. We observed that M-CSF increased the number of donor-derived CD122^+^CD27^+^ NK cell progenitors both in uninfected and infected mice (Fig. 2C).

**Fig. 2.**
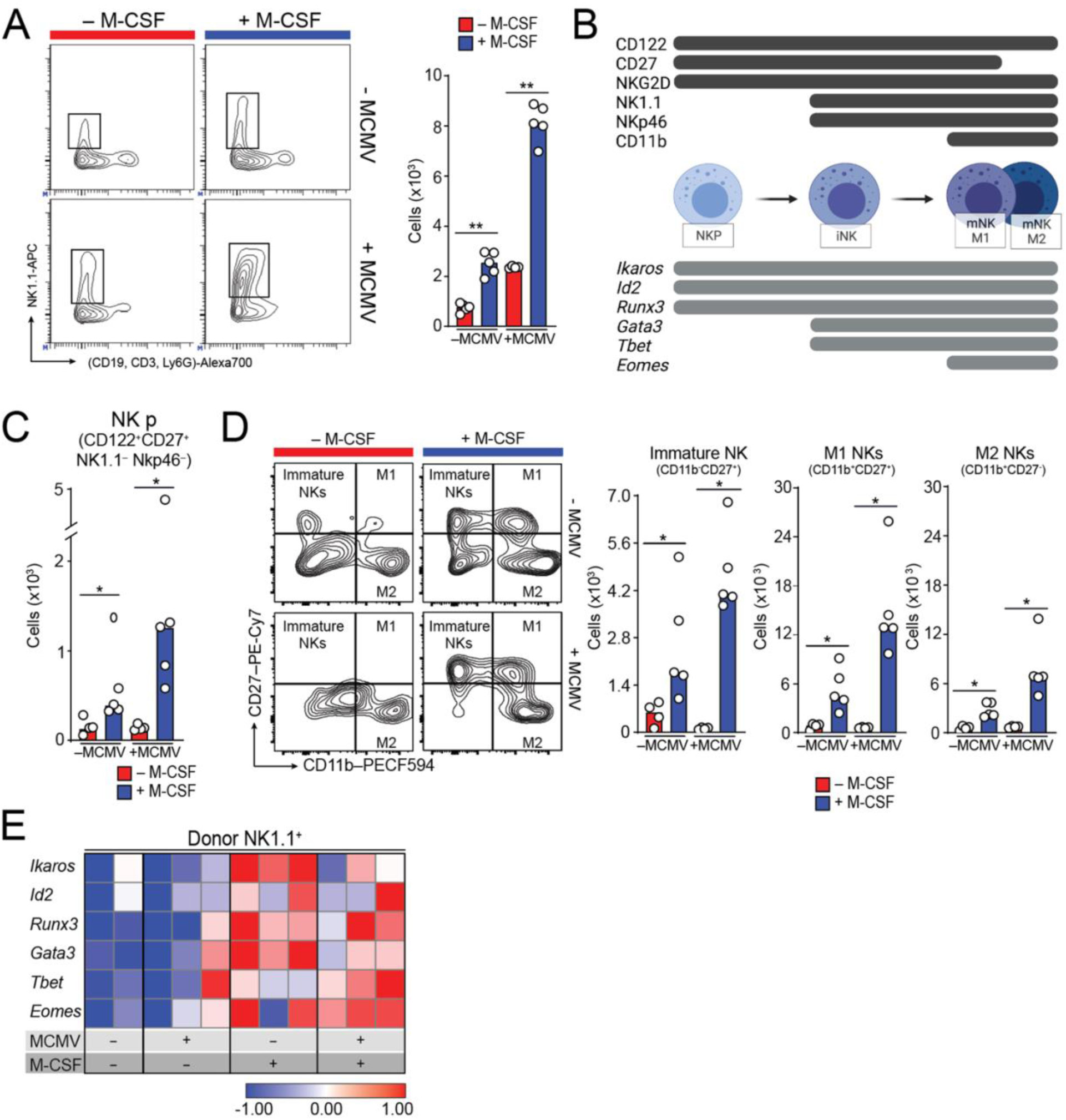
M-CSF treatment increases NK cell production, differentiation, and activation. Experimental set-up as in Fig. 1A. Analysis of spleen NK cell populations. Mice were MCMV- or mock-infected (PBS control) 14 days after HCT (± M-CSF support as indicated in Fig. 1A). Analysis was performed 1.5 days after MCMV or mock infection. (A) FACS examples and median of absolute number of total NK cells (CD19^-^CD3^-^Ly6G^-^NK1.1^+^) are shown. (B) Markers specific to differentiation and maturation stages of NK cells used in this analysis are indicated. (C) Median of absolute number of donor-derived NK progenitor cells (CD122^+^CD27^+^NK1.1^-^Nkp46^-^CD45.1^+^) are displayed. (D) FACS examples and median of absolute numbers of donor-derived immature NK cells, donor-derived M1 (CD11b^+^ CD27^+^) and M2 NK cells (CD11b^+^CD27^-^) are shown. (E) Gene expression analysis of transcription factors expressed by NK cells in FACS-sorted, donor-derived NK1.1^+^ NK cells (definitions of Fig. 2A) by nanofluidic Fluidigm array real-time PCR. \**P* < 0.05, \*\**P* < 0.01 by two-tailed Mann-Whitney *U*-test. All data are representative of two independent experiments.

Consequently, we analyzed the NK cell maturation and differentiation status, which can be distinguished into CD11b^-^CD27^+^ immature, CD11b^+^CD27^+^ mature M1 and CD11b^+^CD27^-^ mature M2 NK cells (Fig. 2B) (*47–49*). M-CSF treatment increased both donor-derived immature and mature M1 and M2 NK cells, particularly in infected mice (Fig. 2D). A smaller increase of progenitor and mature cells was also observed for resident host NK cells (fig. S2B). This was further confirmed by gene expression analysis of stage-specific transcription factors (Fig. 2B). 14 days after M-CSF-supported HCT and after an additional 1.5 days of MCMV or mock infection, spleen NK1.1^+^ cells showed increased expression of the immature NK cell transcription factors *Ikaros*, *Id2*, *Runx3*, *Gata3* and *Tbet* as well as the mature NK cell transcription factor *Eomes* after exposure to MCMV (Fig. 2E). Similar observations were made for host-derived NK cells (fig. S2C). Whereas *Ikaros* and *Gata3* were more strongly induced by M-CSF in uninfected mice, *Tbet* and *Eomes* were preferentially induced after infection (Fig. 2E). Importantly, infection alone was insufficient for the observed inductions.

Together, this indicated that M-CSF lead to an increased number of NK cell progenitors and enhanced their differentiation along the NK cell lineage trajectory.

### NK cells execute M-CSF-derived antiviral immunity

The major antiviral activity of NK cells is mediated by the production of inflammatory cytokines like IFNγ, and perforin-dependent delivery of granzyme B (GrB) into infected cells (*50*). Interestingly, M-CSF treatment increased the number of IFNγ- (Fig. 3A) and GrB-producing NK cells (Fig. 3B) in infected mice in concert with enhanced mRNA levels (*IFNG, GZMB* and *PRF1*) and enriched maturation and activation genes (*CEBPA*, *MITF* and *XCL1*; Fig. 3C, fig. S2D). Consistently, M-CSF induced NK cell accumulation at infectious foci within the liver early after infection culminating in reduced numbers of MCMV-infected cells (Fig. 3D). To determine whether antiviral NK cell activity was required for the protective effect of M-CSF, we depleted NK cells using anti-NK1.1 antibodies in M-CSF-treated and MCMV-infected HCT recipients (Fig. 3E). NK cell-depletion nearly abolished the increased survival of M-CSF-treated mice, demonstrating that a significant part of the protective effect of M-CSF against viral lethality depended on NK cells.

**Fig. 3.**
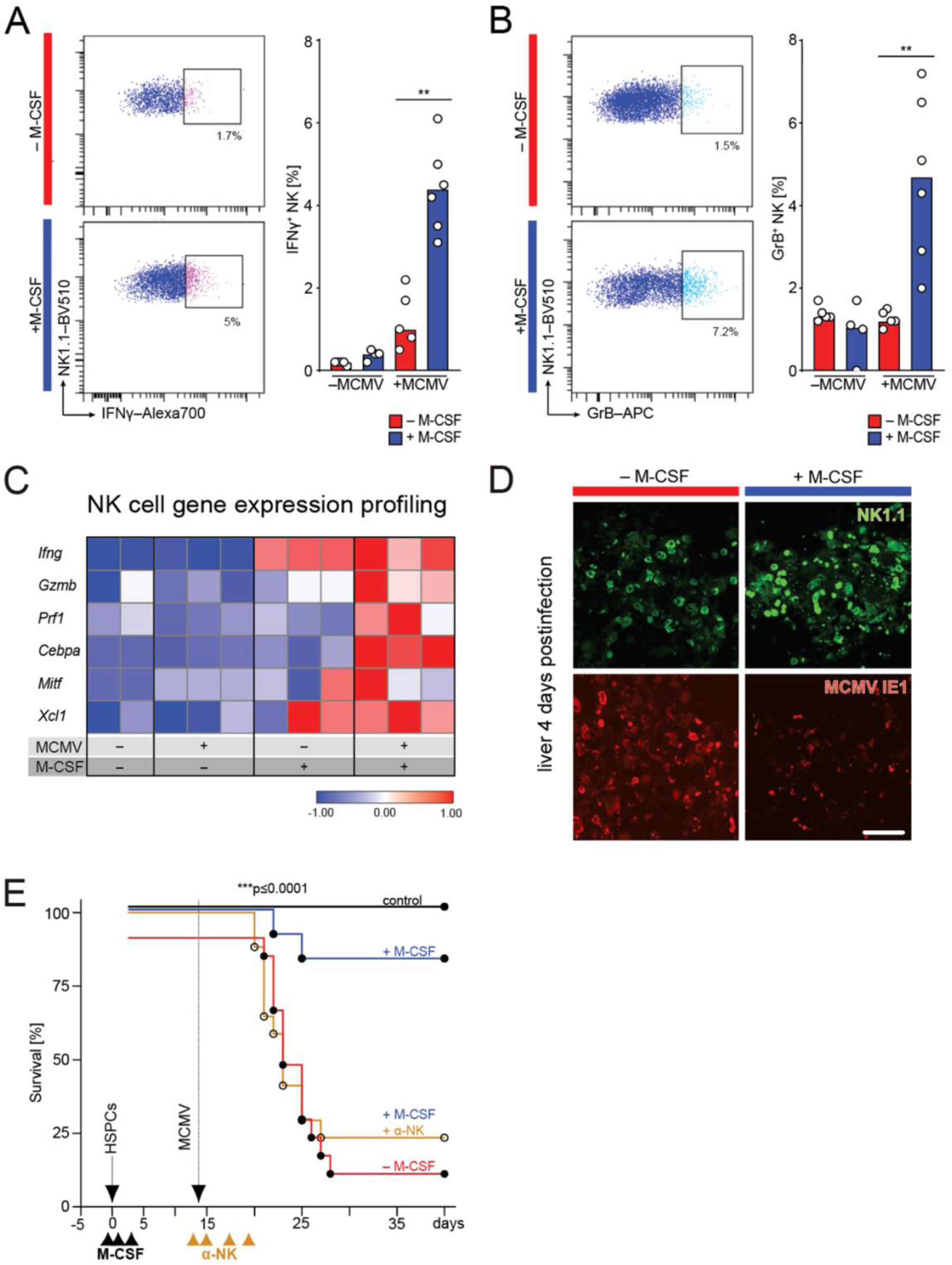
NK cell activity is required for the antiviral effect of M-CSF. Workflow as indicated in Fig. 1A. Analysis was performed 1.5 days (or 4 days in D) after MCMV or mock infection. (A) NK cell activity in the spleen. FACS examples and median of percentage of donor-derived NK1.1^+^ NK cells producing IFNγ. (B) FACS examples and median of percentage of donor-derived NK1.1^+^ NK cells producing GrB. (C) Gene expression analysis of activation and maturation-related factors in FACS-sorted, donor-derived NK1.1^+^ NK cells by RT-qPCR. (D) Immunofluorescence analyses with anti-NK1.1 and anti-MCMV IE1 antibodies in liver of HSPC-transplanted and M-CSF- or control PBS-treated mice 4 days after MCMV infection. (E) Assessment of the requirements for M-CSF-mediated antiviral NK cell response. Survival of PBS control (-M-CSF, n = 15), M-CSF- and control IgG-treated (n = 12), M-CSF and anti-NK1.1-treated (n = 17) or transplanted, uninfected control mice (n = 4). Mice underwent HSPC-transplantation (solid arrow), control PBS or M-CSF-treatment (black arrowheads) and were infected with MCMV <u>(stippled arrow</u>) as shown in Fig.1A-B. Repeated treatment with anti-NK1.1 antibody (or control IgG) was done before and after infection (d-1, d1, d3, d5). \*\**P* < 0.01 by two-tailed Mann-Whitney *U*-test (A, B), \*\*\**P* < 0.0001 by Mantel-Cox test (E). All data are representative of two independent experiments.

### M-CSF-induced myelopoiesis is required for its antiviral effect

Since M-CSF has not been reported to act directly on the NK cell lineage, we investigated whether M-CSF’s effects on the myeloid lineage could indirectly impact on NK cell-mediated antiviral activity. M-CSF treatment can increase donor myelopoiesis in HSPC-transplanted mice (*13, 19*). Accordingly, M-CSF increased donor-derived GMPs, granulocytes, mononuclear phagocytes (Fig. 4A-B), pDCs and cDCs (see fig. S3A-C) two weeks after HCT. To determine whether this was relevant to the antiviral effect of M-CSF, we used complementary loss- and gain-of-function approaches. We injected anti-MCSFR/CD115 antibody 12 days after HCT, which selectively eliminates M-CSF-dependent myeloid cells (*51*). Myeloid cell depletion completely abolished the protective effect of M-CSF treatment in MCMV-infected HCT recipients (Fig. 4C). Affirmatively, these mice showed reduced GMPs, monocytes, cDCs and pDCs 48 hours after anti-CD115 myeloid depletion (Fig. 4D). This indicated that myeloid cells were required for the M-CSF-dependent antiviral activity. For gain-of-function experiments, we transplanted GMPs into mice without M-CSF support 10 days after HCT (Fig. 4A). GMP transplantation resulted in increased survival comparable to M-CSF treatment (Fig. 4E). Together, these experiments demonstrated that the antiviral activity of M-CSF depends upon M-CSF-induced myelopoiesis.

**Fig. 4.**
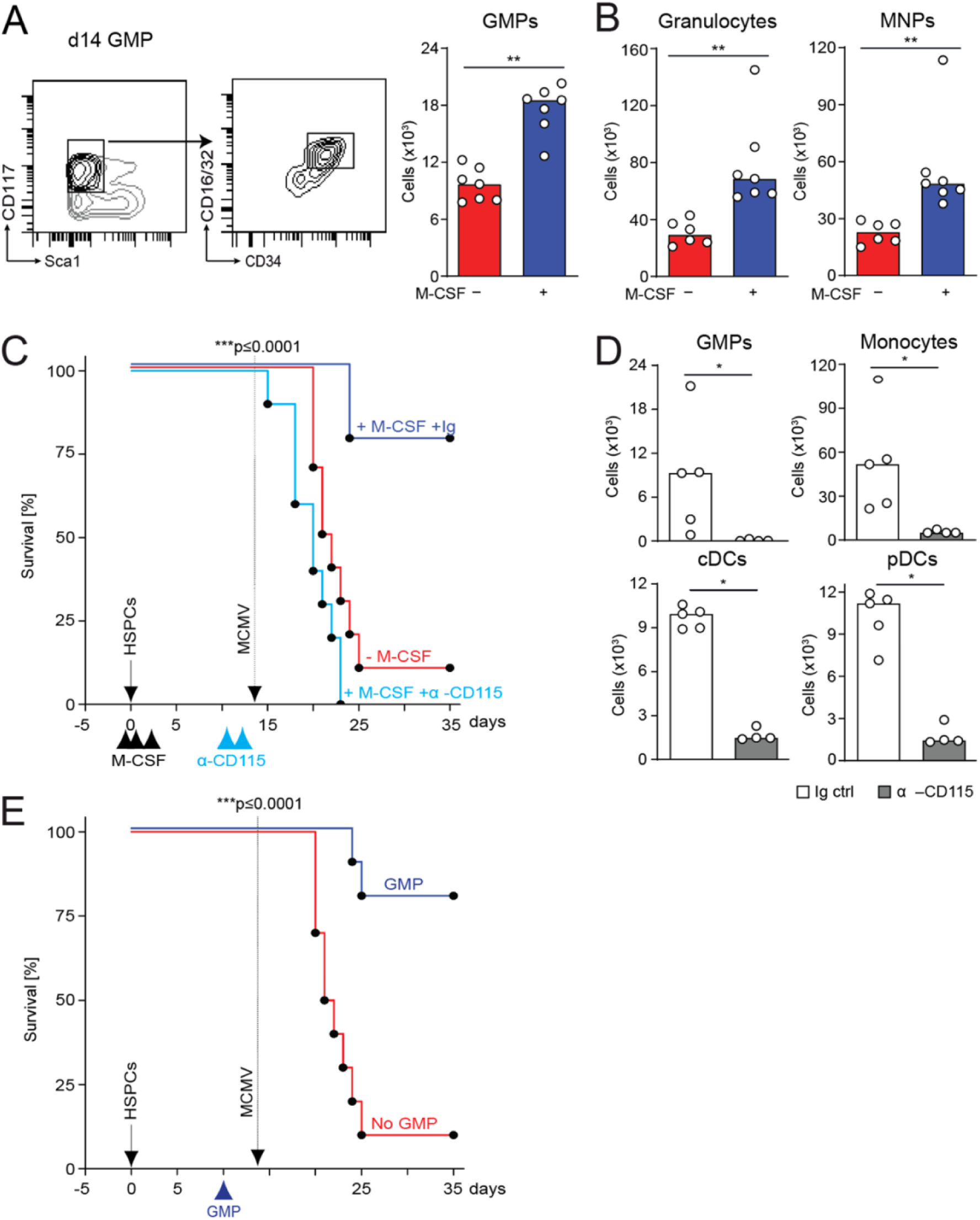
M-CSF-induced myelopoiesis is required for its antiviral effect. (A) Splenic GMPs of control or M-CSF-treated, uninfected mice 14 days after HCT. (B) Splenic granulocytes (Ly6G^+^CD11b^+^) and mononuclear phagocytes (Ly6G^-^CD11b^+^) of control or M-CSF-treated, uninfected recipient mice 14 days after transplantation. (C) Analysis of M-CSF-dependent myeloid cells for its antiviral effect. Survival curve of MCMV-infected and control (n = 10), M-CSF and IgG control Ab-treated (n = 9) or M-CSF and anti-CD115 antibody-treated mice (n = 10). After HCT (solid arrow), vehicle control or M-CSF applied (black arrowheads). Infection with MCMV <u>(stippled arrow</u>) as shown in Fig. 1A-B and treatment twice with anti-CD115 antibody before infection (d-2, d-1). (D) Splenic GMPs, monocytes, cDCs and pDCs of uninfected Ig control- or anti-CD115 antibody-treated recipient mice 48 hours after first treatment. (E) GMP-derived myeloid cells for antiviral activity. GMP transplantation with 50,000 cells on day 10 after HCT. Survival of MCMV-infected control (no GMP, n = 10) or GMP-transplanted mice (GMP, n = 10). Mice underwent HCT (HSPCs) (solid arrow), were infected with MCMV <u>(stippled arrow</u>) as described in Fig. 1A-B and GMP-transplanted 10 days after HCT. \*\*\**P* < 0.0001 by Mantel-Cox test (C, E), \**P* <0.05, \*\**P* < 0.01 by Mann-Whitney *U*-test. All data are representative of two independent experiments.

### M-CSF drives myeloid IL-15 trans-presentation to promote antiviral competence

Since myelopoiesis and NK cell differentiation were required for the antiviral effect of M-CSF, we hypothesized that M-CSF-induced myelopoiesis could indirectly affect NK cell differentiation and antiviral activity. Indeed, anti-CD115-mediated depletion of myeloid cells resulted in reduced immature and mature NK cells (Fig. 5A). To identify myeloid signals that could affect NK cells, we first focused on IL-15, a cytokine paramount for NK cell differentiation and effector functions (*30, 52–54*). IL-15 can be produced and trans-presented by IL15Rα on myeloid cells (*52, 55–57*) including during MCMV infection (*25, 58, 59*). M-CSF treatment resulted in swiftly increased IL-15 mRNA levels in spleens after MCMV infection (Fig. 5B). Since IL-15 signaling requires trans-presentation by the surface molecule IL15Rα (CD215) (*52, 54, 57*), we analyzed the expression levels of IL15Rα in cDCs and monocytes, both capable of stimulating NK cells via IL-15 (*25, 41, 56*). Both mRNA (Fig. 5C) and surface protein analysis (Fig. 5D) revealed that IL15Rα was induced in Ly6C^hi^ monocytes but only weakly in cDCs (Fig. 5C) or Ly6C^lo^ monocytes (Fig. 5D).

**Fig. 5.**
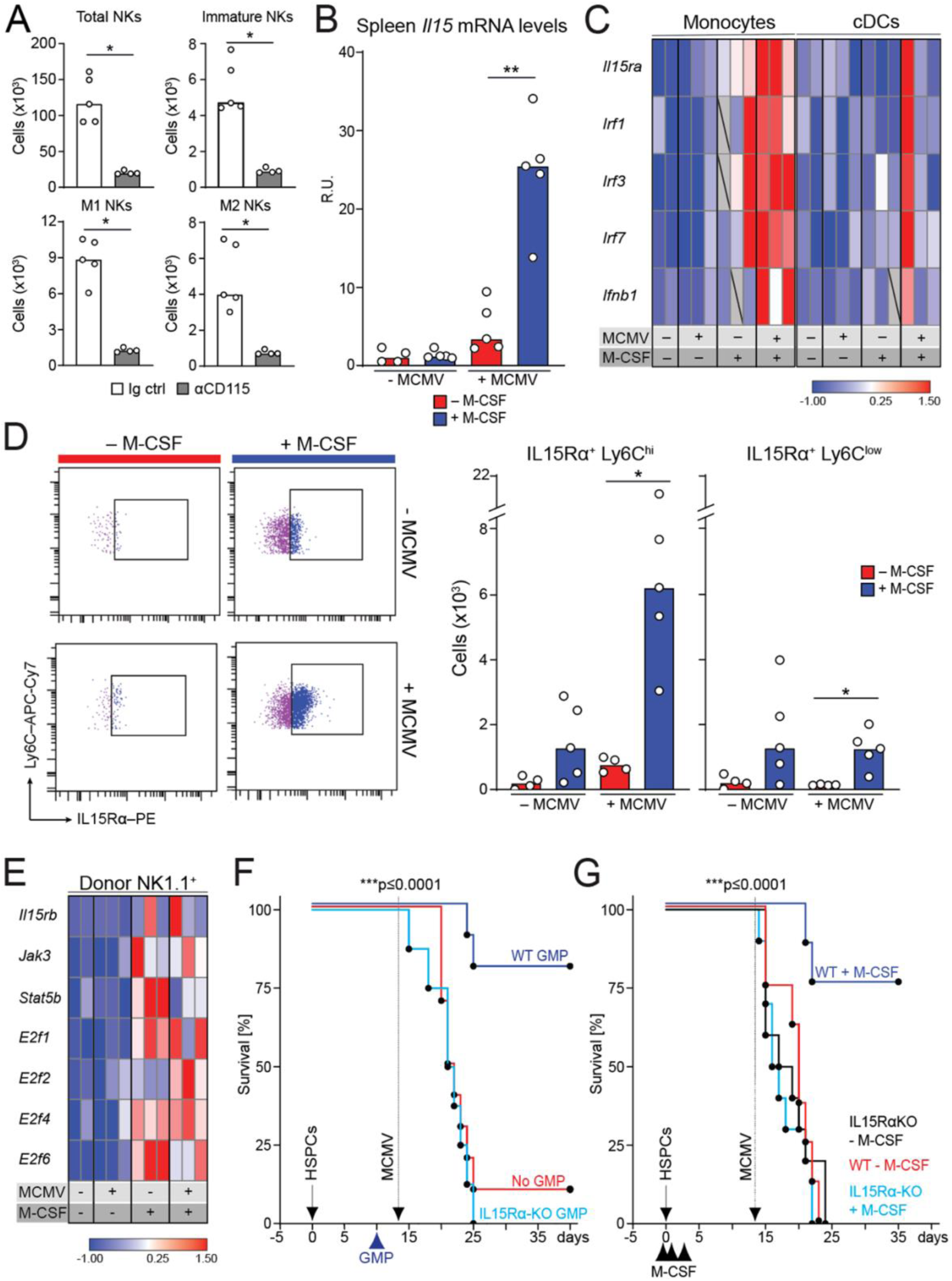
Myeloid IL-15 trans-presentation is required for the antiviral activity of M-CSF. (A) Splenic NK1.1^+^, immature and M1/M2 NK cells of uninfected control or anti-CD115-treated mice two days after depletion and 14 days after HCT and M-CSF treatment. (B) Mice were MCMV- or mock-infected 14 days after HCT. Analysis 1.5 days after infection (B-E). Splenic *Il15* mRNA levels (RT-qPCR). (C) FACS-sorted, donor-derived splenic monocytes and cDCs assessed by Fluidigm. (D) Ly6C^hi^ monocytes (left), splenic IL15Rα-expressing, donor-derived Ly6C^hi^ or Ly6C^low^ monocytes (right). (E) Gene expression analysis of splenic donor-derived NK cells by Fluidigm. (F) 50,000 GMPs transplanted on day 10 after HCT. Survival of MCMV-infected control (no GMP, n = 10), WT GMP (n = 10) or *IL15R*α-KO GMP-transplanted mice (IL15Rα-KO GMP, n = 9). HCT (solid arrow), MCMV infection (stippled) and GMP-transplantation (arrowhead). (G) Survival of WT HCT, control-treated (n = 8), WT HCT, M-CSF-treated (n = 8) or *IL15R*α-KO HCT, control-treated (n = 10) or *IL15R*α-KO HCT, M-CSF-treated mice (n = 10). Mice transplanted with WT control HSPCs or *IL15R*α-KO HSPCs (solid arrow), control or M-CSF treatment (black arrowheads), and MCMV infection (stippled). \**P* < 0.05; \*\**P* < 0.01 by two-tailed Mann-Whitney *U*-test (A-D). \*\*\**P* < 0.0001 by Mantel-Cox test (F-G). All data are representative of two independent experiments.

Next, we analyzed the effect of increased IL-15 signaling from Ly6C^hi^ monocytes on the expression of IL-15 response genes in NK target cells. Like IL-15 signaling, which is engaged once the IL-15/L15Rα complex binds to IL15Rβ on target cells (*25, 52, 55*), M-CSF treatment increased expression of the downstream genes *IL15RB* and of *STATB5*, *JAK3* and *E2F1-6* in NK cells (Fig. 5E). To check whether IL-15-dependent myeloid cell to NK cell-signaling was important for antiviral activity protecting HCT recipients from lethal MCMV infection, we compared *IL15RA*-KO GMPs incapable of trans-presenting IL-15 with WT GMPs. We observed 80% survival in WT GMP-transplanted mice after infection but no survival of *IL15RA*-KO GMP-transplanted mice (50,000 GMPs for each genotype), demonstrating that IL-15 signaling from myeloid cells was required for NK cell support (Fig. 5F). Furthermore, M-CSF treatment resulted in no survival advantage in *IL15RA*-KO HCT recipient mice and was comparable to untreated WT HCT mice (Fig. 5G), indicating that IL-15 signaling was acting downstream of M-CSF.

Together, our data demonstrate that myeloid-derived IL-15 signaling was required for the antiviral effect derived from M-CSF-induced myelopoiesis.

### M-CSF-induced I-IFN production stimulates IL-15-dependent antiviral immunity

I-IFNs contribute to the early antiviral immune response preceding NK cell activation (*22, 25, 60–62*), and thus, may constitute a rapid response mechanism that could prevent fatal viremia during leukopenia after HCT. MCMV infection was shown to increase *IFNB1* mRNA in the spleen (*26, 27*). Consistently, we found enhanced *IFNB1* mRNA levels in the spleen swiftly after MCMV infection, which were augmented with M-CSF treatment (Fig. 6A). During MCMV infection, I-IFNs are predominantly produced by pDCs. We observed that pDC numbers (Fig. 6B) and I-IFN-producing pDCs (Fig. 6C) were increased in the spleen of M-CSF-treated mice 14 days after HCT, particularly after MCMV infection. Monocytes showed a strongly increased expression of *IFNB1* and upstream transcription factors of the IRF family (Fig. 5C). Together, this supported the notion that M-CSF treatment increased I-IFN production during MCMV infection of HCT recipients by promoting a faster reconstitution of monocytes and pDCs. This also agrees with the observation that M-CSF-driven myelopoiesis can also stimulate pDC development (*63*). The observed effects of both loss- and gain-of-function experiments targeted at myeloid cells (Fig. 4C-D) or transplantation of GMPs, which also give rise to pDCs, thus support the notion of I-IFNs also contributing to antiviral immunity upon M-CSF administration after HCT.

**Fig. 6.**
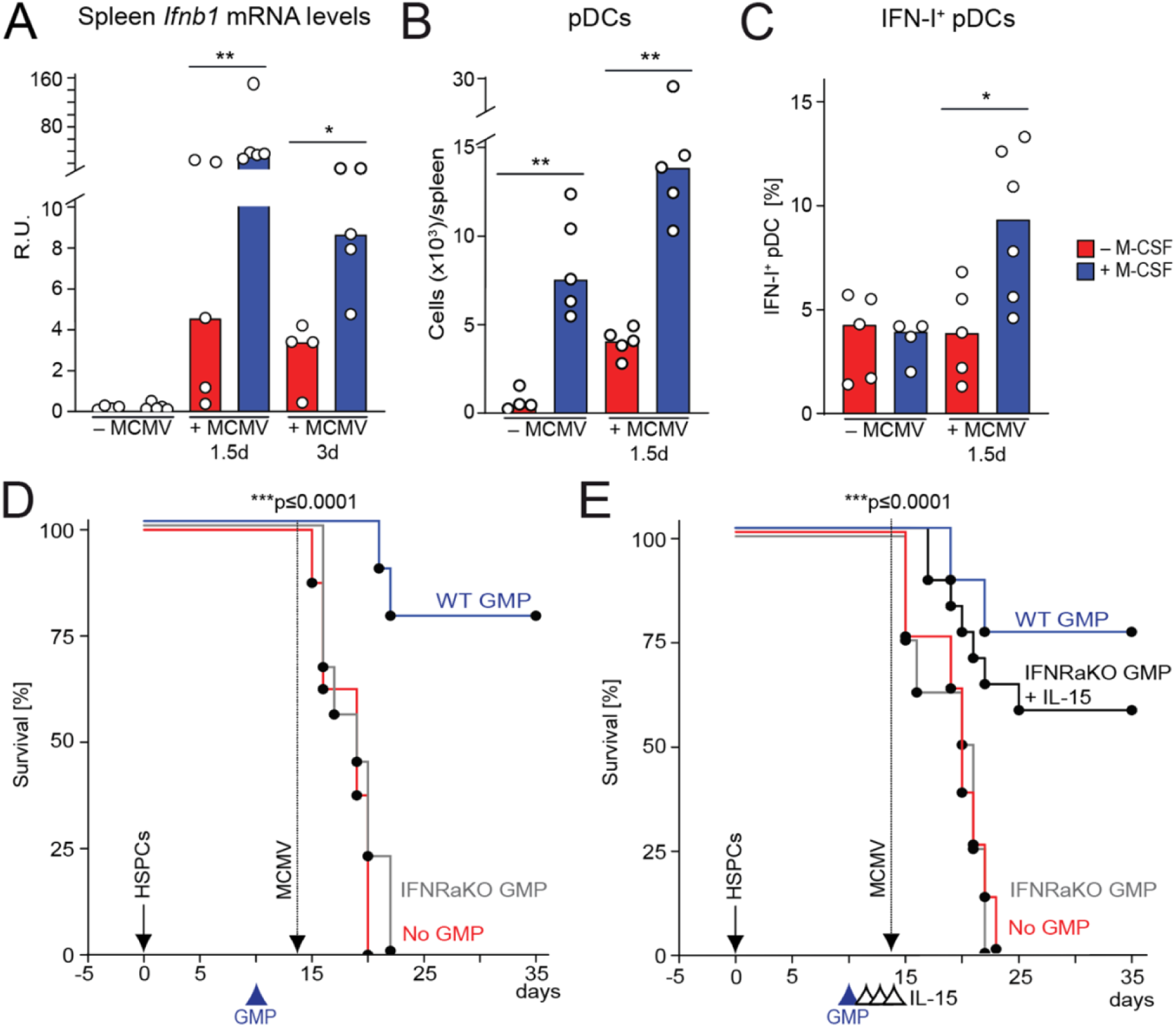
M-CSF-induced I-IFN production stimulates IL-15-dependent antiviral effects. MCMV- or mock-infection 14 days after HCT. Analysis performed 1.5 days (or 3 days in A) after. (A) Splenic *Ifnb1* mRNA levels of control or M-CSF-treated mice (RT-qPCR). (B) Donor-derived splenic Lin^-^CD11c^lo^BST2^hi^ pDCs. (C) %IFN-β^+^ pDCs. (D) Survival of MCMV-infected, no GMP control (n = 8), WT GMP (n = 9) or *Ifnar1*-KO GMP-transplanted mice (n= 9). HCT (solid arrow), MCMV infection (stippled) and GMP-transplantation 10 days after HCT (arrowhead). *P* < 0.0001 by Mantel-Cox test. (E) Survival of MCMV-infected, no GMP control (n = 8), WT GMP (n = 8) or *Ifnar1*-KO GMP-transplanted mice without (n = 8) or with IL-15 rescue treatment (n = 16). HCT (solid arrow), MCMV infection (stippled), GMP transplantation 10 days after HCT (blue arrowhead) and treatment with 0.5 µg IL-15 or control on days 12, 13 and 14 (black arrowheads). \*\**P* < 0.01, \**P* < 0.05 by Mann-Whitney *U*-test (A-D). \*\*\**P* < 0.0001 by Mantel-Cox test (E-F). All data are representative of two independent experiments.

Beyond its direct antiviral effects on infected cells, I-IFNs can also indirectly affect the antiviral immune response by activating NK cells or by stimulating IL-15 production in myeloid cells (*25, 30, 62*). To investigate the relative importance of I-IFNs on myeloid cells, we injected *IFNAR1*-KO or WT GMPs at day 10 after HCT. *IFNAR1* deficiency abolished the protective effect of GMP transplantation (Fig. 6D), indicating that I-IFN stimulation of myeloid cells was required for their antiviral effect. IL-15 treatment prior to infection could partially restore the deficiency of *IFNAR1*-KO GMPs (Fig. 6E), indicating the importance of I-IFN induction of IL-15 production in myeloid cells.

Together, this suggested that the antiviral activity of I-IFNs was mainly due to its effect on the identified myeloid and NK cell differentiation program rather than a direct effect on infected cells.

### M-CSF recapitulates its effects in human G-CSF-mobilized PBMCs

To determine whether M-CSF could affect myeloid and NK cell differentiation in human HSPCs, we assayed its impact on myelopoiesis, IL15Rα expression, NK cell numbers and functional competence in HSPC-enriched PBMCs from G-CSF-mobilized stem cell donors (G-PBMCs). *In vitro* differentiation from G-PBMCs was established in the presence of stem cell factor (SCF) alone or in combination with the multi-lineage myeloid cytokine IL-3 or with M-CSF, respectively (protocol fig. S4A and the gating strategy used fig. S4B-E). Both IL-3 and particularly M-CSF fostered myelopoiesis, yielding increasing proportions of cells with monocytic and/or macrophage morphology (Fig. 7A). This observation was confirmed by flow cytometry, where we found faster and stronger reduction of CD34^+^ progenitors (Fig. 7B-C) and a concomitant increase of CD11b^+^ myeloid cells for M-CSF conditions (Fig. 8A-B). Consistent with a faster myeloid commitment in the presence of M-CSF, we also found increased numbers of GMPs (Fig. 7D, 8C), in particular HLA-DR^+^ mature GMPs (Fig. 8D) (*64, 65*). The enhanced and accelerated frequencies of CD11b^+^ myeloid cells after M-CSF treatment were mainly due to CD11b^+^CD66b^-^ monocytic cells (Fig. 8E) rather than CD11b^+^CD66b^+^ granulocytic cells (i.e., neutrophils; data not shown). Enhanced monocytic differentiation was further confirmed by accelerated and increased CD14^+^ monocyte generation at days 5 and 9 after M-CSF treatment (Fig. 8F). Whereas monocytes isolated from freshly isolated G-PBMCs consisted mainly of CD14^+^CD16^-^ classical monocytes (CMs) (Fig. 7E, 8G), at day 9 after M-CSF treatment we observed nearly exclusively CD14^+^CD16^+^ intermediate monocytes (IMs), with non-classical CD14^-^CD16^+^ monocytes (NCMs) (*66*) remaining low under both conditions (Fig. 7E, 8G).

**Fig. 7.**
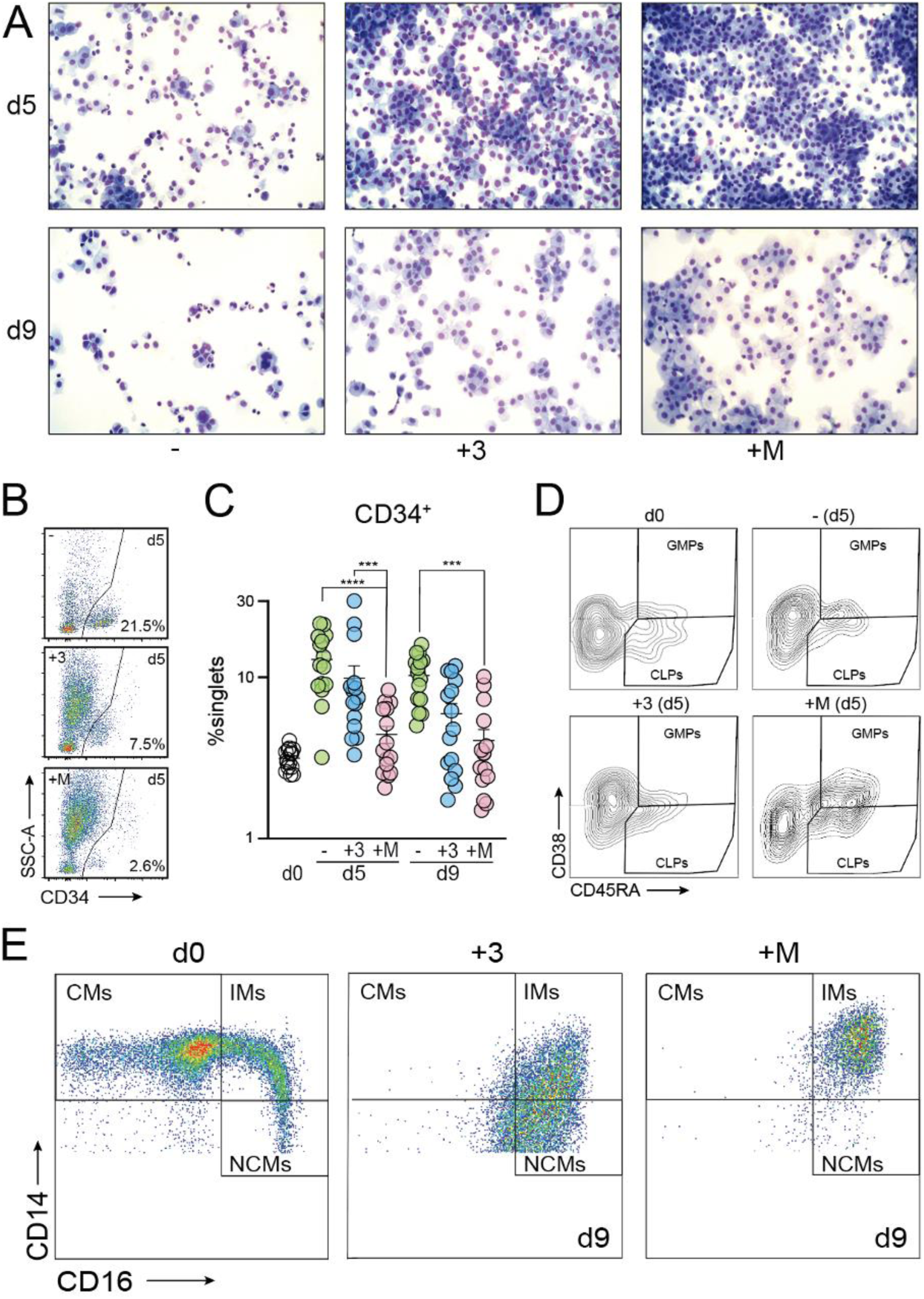
M-CSF supports terminal differentiation of IMs in human G-PBMCs. (A) Cytospins at days 5 or 9 after *in vitro* differentiation without myelopoiesis-inducing cytokines (-), or with IL-3 (+3) or M-CSF (+M) (modified Giemsa). (B) Contour plots five days after *in vitro* cytokine treatment without myelopoiesis-inducing cytokines (-, top row), with IL-3 (+3, middle row) or with M-CSF (+M, bottom row) indicating the frequency of CD34^+^ HSPCs as compared to the total number of live single cells analyzed. (C) Frequency of CD34^+^ HSPCs at seeding (d0), or after *in vitro* cytokine treatment without myelopoiesis-inducing cytokines (-, green circle), with IL-3 (+3, blue circle) or with M-CSF (+M, salmon circle). (D) Contour plots of HSPC populations comprising of CLPs (CD34^+^CD45RA^+^CD38^-^) or GMPs (CD34^+^CD45RA^+^CD38^+^) upon selection from G-PBMCs (d0) or after *in vitro* cytokine treatment at day 5: without myelopoiesis-inducing cytokines (-) vs. IL-3 (+3) vs. M-CSF (+M). (E) Seeding (d0): monocytes SSC-A^low^CD14^+^ (CMs by CD16/CD14 staining, left); differentiation: IMs almost exclusively with M-CSF (+M, right) vs. 2/3 IMs and 1/3 NCMs with IL-3 (+3, middle) at day 9. Representative pseudocolor plots. The data are illustrated as mean ± SEM. A ratio-paired *t*-test was used. \*\*\**P* < 0.001, \*\*\*\**P* < 0.0001. All data are representative of five independent experiments.

**Fig. 8.**
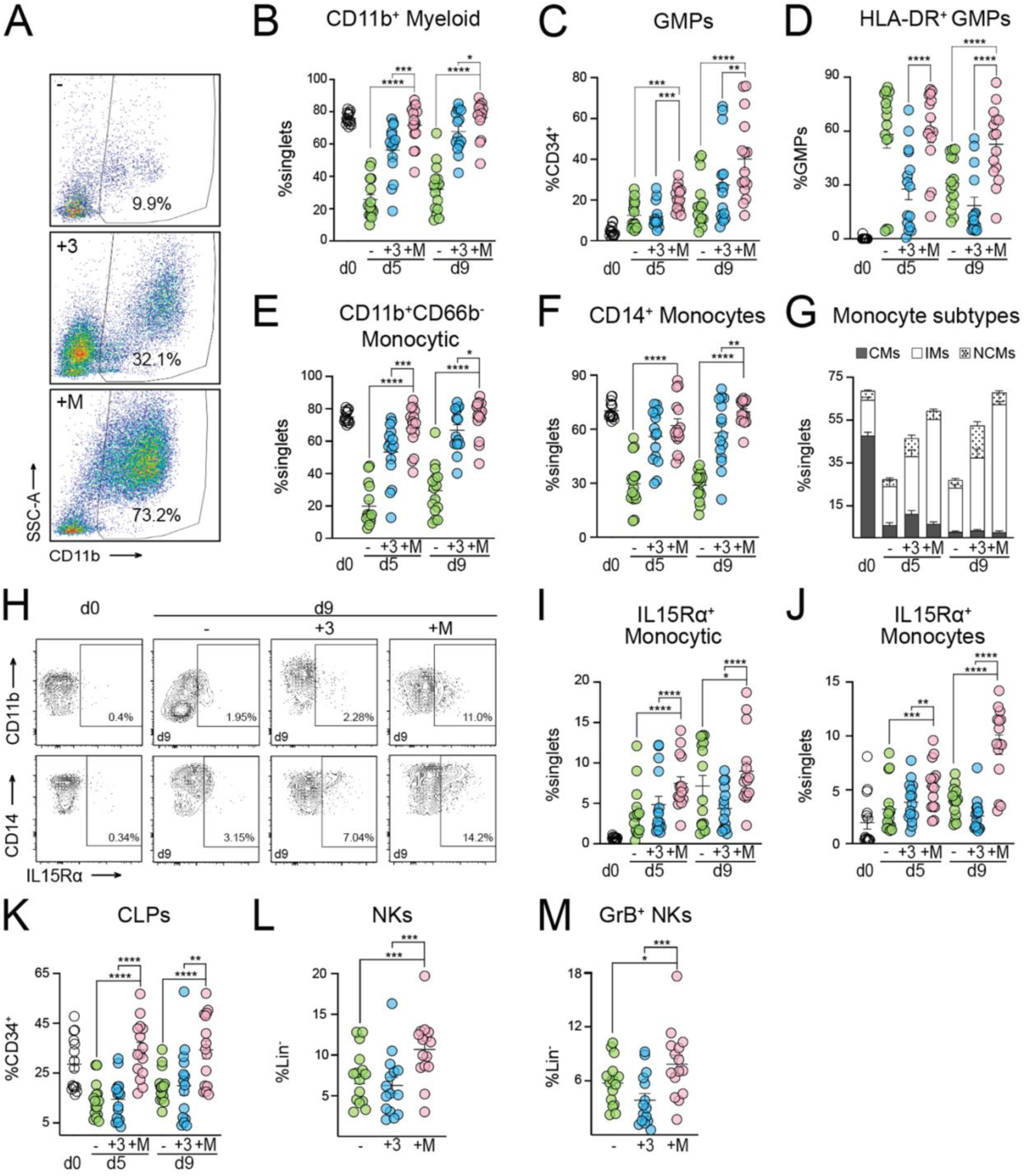
M-CSF-driven myelopoiesis induces IL15Rα expression on monocytes and supports NK cell viability and cytokine competence in human G-PBMCs. (A) Contour plots of G-PBMCs in G-CSF-mobilized donors five days after *in vitro* cytokine treatment without myelopoiesis-inducing cytokines (-, top row), with IL-3 (+3, middle row) or with M-CSF (+M, bottom row). (B) Quantification of CD11b^+^ cells. (C) Frequency of GMPs at seeding (d0, empty circle) or without myelopoiesis-inducing cytokine treatment (-, green circle), with IL-3 (+3, blue circle) or with M-CSF (+M, salmon circle). (D) Frequency of mature GMPs (HLA-DR^+^) at seeding or -, or +3 or +M. (E) M-CSF-driven myelopoiesis. (F) M-CSF-driven monopoiesis. (G) M-CSF stimulates intermediate monocytes (IMs); CMs: classical monocytes, solid bars, IMs: white bars, NCMs: non-classical monocytes, dotted bars. (H) M-CSF drives IL15Rα expression. Contour plots of IL15Rα expression on CD11b^+^ and CD14^+^ cells. (I) Quantification of (H) on CD11b^+^CD66b^-^ monocytic cells. (J) Quantification of (H) on CD14^+^ monocytes. (K) Quantification of CLPs. (L) M-CSF supports cytokine-competent NK cells (NKs, SSC-A^low^Lin^-^ CD56^+^CD16^+^) compared to - and +3. (M) M-CSF treatment enhances Granzyme B (GrB) production in NKs compared to IL-3 and -. The data are illustrated as mean ± SEM. A ratio-paired *t*-test was used. \**P* < 0.1, \*\**P* < 0.05, \*\*\**P* < 0.01, \*\*\*\**P* < 0.001. All data are representative of five independent experiments.

Together, these data indicated that M-CSF also resulted in increased monopoiesis in human HSPCs with a particular enrichment in IMs.

As observed in murine cells, M-CSF treatment also resulted in enhanced IL15Rα expression on CD11b^+^ myeloid cells and CD14^+^ monocytes (Fig. 8H) with increasing levels during differentiation on CD11b^+^CD66b^-^ monocytic cells or CD14^+^ monocytes (Fig. 8I-J). Together, this indicated that M-CSF treatment also enhanced IL-15 presentation on human monocytes. Finally, we further queried the effect of M-CSF-driven myelopoiesis and IL15Rα signaling in human G-PBMCs on functional NK cell differentiation. We first analyzed CLPs, which encompass NK cell progenitors (*67*). Interestingly, CLPs were enriched in M-CSF-treated G- PBMCs, both at days 5 and 9 (Fig. 8K). Although the culture regime lacked exogenous IL-2, IL-15 or IL-21 and thus was not ideal for NK cell differentiation and survival, M-CSF-driven myelopoiesis resulted in significantly more NK cells (NKs, SSC-A^low^Lin^-^CD56^+^CD16^+^) at day 9 of culture (Fig. 8L). In line with the findings in murine cells, M-CSF treatment also increased the numbers of GrB-expressing NKs (Fig. 8M) significantly on day 9 of culture compared to IL-3-driven myelopoiesis, indicating that M-CSF treatment also stimulated functionally competent human NK cell production.

Together, these findings indicated that the coordinated myeloid-driven NK cell differentiation and activation program initiated by M-CSF-mediated myelopoiesis in mice was translatable to the human context and was thus directly relevant for clinical conditions of HCT.

### No adverse events of M-CSF after allogeneic HCT

In hematooncology several indications require allogeneic HCT. To date, there are conflicting data concerning the effect of M-CSF on long-term engraftment and GvHD following allogeneic HCT (*68–70*). Hence, we used an allogeneic HCT model to address these points (fig. S5A). Following allogeneic HCT and assessment according to previous reports (*68*), we did not find any differences in the frequency of C57BL/6j CD45.1^+^ donor HSPC-derived CD11b^+^F4/80^+^ monocytes or inflammatory Ly6C^HI^ monocytes after M-CSF treatment (fig. S5B-C).

Furthermore, GvHD scoring (*71*) showed no statistical difference between mice treated with M-CSF or PBS, going to the lowest possible score as early as 20 days following allogeneic HCT for both conditions (fig. S5D). All mice survived M-CSF treatment and vehicle control after allogeneic HCT. We further showed that tri-lineage engraftment in the peripheral blood was not affected by M-CSF treatment at 4 and 12 weeks following allogeneic HCT (fig. S5E). Merely any residual recipient BALB/c CD45.2^+^ cells were found (“alloHSCs” in fig. S5B), reflecting full bone marrow (BM) engraftment of CD45.1^+^ donor cells, which we confirmed at 12 weeks (fig. S5F). These CD45.1^+^ donor cells were unaffected by the M-CSF treatment concerning long-term engrafting HSPCs (KSL Flt3^-^CD150^+^CD48^-^) and GMPs alike in the BM 12 weeks after allogeneic HCT (fig. S5G).

Together, our data reveal no contraindication for the short-term treatment with M-CSF following allogeneic HCT, suggesting that it should be a safe and feasible cytokine to promote antiviral activity in standard protocols of allogeneic HCT.

## DISCUSSION

In this study we have identified the previously unknown protective effects of M-CSF-induced myelopoiesis against viral infection during the vulnerable leukopenic phase after HCT. We identified a coordinated differentiation program between myeloid and NK cells that plays a major role in reconstituting protection against viral infection and assigns a critical role to M-CSF-induced myelopoiesis in participating in antiviral immunity. Immunocompromised individuals are prone to opportunistic infections including CMV viremia, but also to infection-induced morbidity and mortality (*1*). Here, we used a murine model of immunosuppression after HCT to investigate the protective antiviral effects of M-CSF-induced myelopoiesis preceding MCMV infection. HCT is an important major therapeutic strategy that involves a conditioning therapy by which the recipient’s hematopoietic system is immunosuppressed to foster engraftment of donor HSPCs. Patients encounter severe immunodeficiency after HCT that leaves them highly vulnerable to opportunistic bacterial, fungal, and viral infection before the donor’s hematopoietic system is sufficiently reconstituted. Although improvements have been made in prophylaxis and management, viral infection, and reactivation, such as CMV, still contribute significantly to morbidity and mortality after allogeneic HCT (*3, 72, 73*). Unfortunately, available antiviral drugs are associated with numerous adverse events (*2, 5*). For example, ganciclovir severely compromises myelopoiesis, and thus further aggravates susceptibility to secondary infections (*74–76*) and enhances risk to secondary malignancy (*77*). Although progress has been made with the introduction of letermovir as non-toxic antiviral agent, it might select for virus variants and virus breakthrough infections as well as late reactivation once cessation of prophylaxis occurs (*7*). Furthermore, as an agent targeting viral terminase complex it is limited to be used against CMV. Adoptive transfer protocols of lymphoid progenitors also have been proposed as a therapeutic strategy in refractory or high-risk cases (*8*). Cell therapy approaches, however, require complex logistics, which limits their availability and leads to high costs. Given the remaining clinical need for both acute and prophylactic antiviral treatments, the application of M-CSF may represent an attractive, cost-effective, and broadly applicable antiviral approach.

Several properties of M-CSF make it an ideal candidate for accelerating immunocompetence recovery in HCT recipients and present key advantages over other myeloid cytokines used in clinical practice. We showed before that M-CSF directly engages HSPCs and thus intervenes at the earliest point of the differentiation hierarchy to initiate the production of innate immune cells (*13, 18, 19*). M-CSF prophylaxis could therefore shorten the time of immune system reconstitution to reduce the risk of infections. Other cytokines, in particular G-CSF, are also used to stimulate immune functionality. However, in contrast to M-CSF, G-CSF can only act on already existing mature or late myeloid progenitor cells to activate their functional competence. Since these cells will only develop weeks after HCT, G-CSF will be ineffective in the early phase after HCT. By acting at the earliest point of the hematopoietic differentiation hierarchy, M-CSF can stimulate myelopoiesis swiftly after conditioning therapy. Consistent with this, we showed previously that M-CSF but not G-CSF can stimulate the increased production of myeloid cells from HSPCs and protect from bacterial and fungal infections (*19*). Importantly, M-CSF-induced myelopoiesis neither compromises stem cell numbers or activity (*18*), nor comes at the expense of the generation of other blood cell lineages like platelets that are important for restoring blood clotting activity (*19*). Here, we report an additional advantage of M-CSF treatment by promoting rapid reconstitution of antiviral activity and protection from viral infection through a multistep myeloid and NK cell differentiation program. A significant advantage of the early action of M-CSF on HSPCs appears to be the stimulation of a combination of innate immune cells that are required to combat pathogens. Whereas G-CSF only stimulates granulocytes and their direct progenitors, M-CSF stimulates the production of i) granulocytes, mediating cytotoxic bacterial killing, ii) monocytes and macrophages, capable of pathogen control by phagocytosis and reactive oxygen production, and iii) dendritic cells with the strongest antigen presentation activity that alerts the adaptive immune system. In this study, we now show that M-CSF also induced I-IFN-producing pDCs and indirectly stimulated NK cell differentiation and activation through induction of IL-15-producing monocytic cells, which together mediated strong antiviral activity.

Clinical protocols of HCT often involve allogeneic HCT and thus harbor the risk of GvHD. Interestingly, M-CSF treatment ameliorated GvHD after allogeneic HCT in a murine model (*70*), where M-CSF was applied at the time of transplantation similar to the protocol used in this study. A seemingly conflicting study, showing increased GvHD after M-CSF administration (*68*), applied M-CSF at a much later point in time. In this study, M-CSF was applied after at least two weeks, where it probably acted on infiltrating monocytes and macrophages in GvHD-affected tissue sites. In line with a beneficial role of M-CSF in GvHD, clinical data from 54 patients treated with M-CSF after allogeneic HCT revealed no difference in the frequency of chronic GvHD, but severe GvHD was rather attenuated by M-CSF application compared to control groups (*69*). Accordingly, we observed no detrimental effect of M-CSF in allogeneic HCT mice (fig. S5B-C). Together, this indicates that M-CSF prophylaxis can boost antiviral immunity following HCT and may bring about the additional benefit of reducing the occurrence and severity of GvHD. The M-CSF prophylaxis described by us targets NK cells and pDCs, whose protective functions during CMV infection are well described (*22, 78*). This is important for a fast antiviral response under immunosuppressed and leukopenic conditions since an antiviral T cell response is not required and cannot be mounted. This is particularly beneficial for standard protocols of HCT, which are commonly T cell depleted. Under these circumstances, the development of engrafted T cells arising from donor HSPCs occurs much later than viral reactivation during immunosuppressive leukopenia.

The effect on NK cells described in this study is mediated by M-CSF-induced myelopoiesis, in particular monocytes. The role of monocytes and macrophages in CMV infection is multifaceted. On the one hand, they can be target cells for MCMV infection (*79–81*), thus serving as vehicles of CMV dissemination (*32, 82*). On the other hand, the observation that macrophage depletion increased MCMV burden (*79*), also support a protective role during CMV infection. This ambiguity might be dependent on the context of infection or on the specific monocyte subpopulation. Whereas Ly6C^-^CX3CR1^hi^ patrolling monocytes are involved in dissemination (*32*), Ly6C^+^CCR2^+^ inflammatory monocytes can engage antiviral responses in early infection via direct or indirect mechanisms (*33, 34, 38, 80, 83*). Ly6C^+^CCR2^+^ inflammatory monocytes could initiate differentiation of memory CD8^+^ T and NK cells into antimicrobial effector cells (*33*) or showed direct iNOS-mediated antiviral effects (*34*). I-IFN signaling is also important for recruitment of CCR2^+^ inflammatory monocytes via MCP-1/CCL2 (*83*). Thus, mice deficient for MCP-1 or CCR2 showed a reduced accumulation of monocyte-derived macrophages and NK cells in liver, increased viral titers, widespread virus-induced liver pathology and reduced survival (*80, 84*). Previously, it was shown that CD11c^hi^ DC-derived IL-15 promoted NK cell priming (*56*) and that inflammatory monocyte-derived IL-15 could stimulate NK cell differentiation (*33*). In the immunosuppressed settings investigated here, Ly6C^hi^ monocytes appeared to be more important than DCs for IL-15 presentation, since they expressed higher levels of IL15Rα required for IL-15 cross-presentation to NK cells (*56*). Consistent with the synergistic role of IL-15 and I-IFNs for NK cell activation (*30, 56*), we observed that both *IL15RA*- and *IFNAR1*-deficiency in GMP-derived myeloid cells abolished their protective effect against MCMV, whereas ectopic IL-15 could rescue *IFNAR1*-deficiency. This suggested that IL-15 induction in monocytes required I-IFNs that were mainly produced by M-CSF-induced pDCs. Together, our experiments revealed the surprising capacity of M-CSF to initiate a fully synchronized differentiation program and cytokine mediated crosstalk between different myeloid and NK cell lineages to provide effective antiviral prophylaxis during leukopenia following HCT-mediated immunosuppression.

However, further studies are needed to evaluate the clinical employability of M-CSF after HCT as a prophylaxis of CMV infection in humans. For this, phase I/II clinical trials will be needed to evaluate the addition of M-CSF to the currently licensed cytokine treatment options comprising of G-CSF and GM-CSF.

## MATERIALS AND METHODS

### Study design

The experiments in this study were design to examine the relevance of M-CSF to support myeloid reconstitution and myeloid-driven support of antiviral competence of NK cells against dormant viruses such as Herpesviridae. For this reason, we chose CMV since it is highly relevant in the human context causing high morbidity and mortality after HCT. We sought to investigate the mechanism of M-CSF-driven antiviral protection. WT or gain-of-function/loss-of-function approaches were used in C57BL6 mice. The animals were used to assess the *in vivo* impact of M-CSF on viral load, histological features of CMV pathology, impact on myeloid and NK cell differentiation, or overall survival by RNA profiles and flow cytometry and using antibody-depletion approaches Allogeneic transplantations of C57BL6/j mice stem cells into BALB/c recipient mice were performed to demonstrate safety of M-CSF administration and its effect on myeloid reconstitution in an allogeneic HCT model. All mouse experiments were performed under specific pathogen-free conditions in accordance with institutional and national guidelines under permit numbers APAFIS #17258-2018102318448168-v5 and APAFIS #36188-2022032912082580-v6 monitored daily for signs of morbidity. To demonstrate translatability of the M-CSF-induced mechanisms in the G-CSF-mobilized PBMCs from human stem cell donors were obtained from leukapheresis samples from the Department of Transfusion Medicine of the TU Dresden. The use of human samples was approved by the ethical review committee of the TU Dresden (approval no. EK477112016 and EK393092016) and all human research conformed to the Declaration of Helsinki. Informed consent was obtained from all participants.

### Mice and *in vivo* treatments

For reconstitution 3,000 c-Kit/CD117^+^Sca1^+^Lin^-^ HSPCs, isolated using a lineage depletion kit (Miltenyi Biotec) and FACS sorting from 6–8-week CD45.1^+^ bone marrow, were injected with 150,000 cKit^-^Ter119^+^ CD45.2^+^ carrier cells (Miltenyi Biotec) and murine (baculovirus expressed) or human recombinant M-CSF (Chiron/Novartis) in 200 µL PBS retroorbitally into lethally irradiated (160 kV, 25 mA, 6.9 Gy) 8-14 weeks sex-matched CD45.2^+^ mice as described previously (*13, 18*). Myeloid or NK cells were depleted by multiple intraperitoneal injections of 100 µg of rat anti-CD115 (*51*), anti-NK1.1 mAb (*60*) or control IgG in PBS before MCMV infection as indicated. 50,000 granulocyte-monocyte progenitors (GMPs) (Lin^-^CD117^+^Sca^-^1^-^ CD34^+^CD16/32^+^) from WT or *IL15R*α-KO or *IFNAR1*-KO mice were FACS sorted and injected on day 10 after HCT.

For allogeneic HCT, BALB/c CD45.2^+^ recipient and C57BL/6j CD45.1^+^ donor mice were used. In brief, BALB/c CD45.2^+^ recipient mice were irradiated with 5 Gy, followed by allogeneic HCT after 24 hours. Imminently before (one hour) or shortly after (five and 20 hours) allogeneic HCT with 2 x 10^5^ Lin^-^ HSPCs from C57BL76j CD45.1^+^ donors, the mice received PBS or 10 µg of baculoviral expressed human M-CSF. Following alloHCT, scoring for graft-versus-host-disease (GvHD) was performed according to Lai *et al*. (*71*) on days five, ten, 13, 15, 20 and 30, as well as donor HSPC-derived blood cells were ascertained on day 30 in accordance with Alexander *et al*. (*68*).

### MCMV infection, viral loads and histopathology

Two weeks after HCT, mice were injected intraperitoneally with 5,000 PFU MCMV K181 v70 in 200 µL PBS. Viral loads were measured by quantitative reverse transcription polymerase chain reaction (RT-qPCR) of *Ie1* mRNA (*25*) extracted from frozen tissues 36-40 hours (1.5 days) or 72 hours as reported previously (*60*). Paraformaldehyde-fixed (4%), paraffin-embedded and hematoxylin and eosin (H&E)-stained liver sections were scored by a trained veterinary pathologist blinded to sample identity for indicated parameters.

### Human hematopoietic stem and progenitor cell differentiation

Human G-CSF-mobilized HSPCs were obtained from leukapheresis samples from the Department of Transfusion Medicine of the TU Dresden. On the day of donation, a Ficoll-density gradient centrifugation step was performed as described previously (*85*) to isolate the peripheral blood mononuclear cell (PBMC) layer containing mainly mononuclear cells, T cells, NK cells, HSPCs and low-density granulocytes. To evaluate the effect of M-CSF on selective co-cultures between mononuclear cells, NK cells and HSPCs, T cells and low-density granulocytes were depleted with anti-CD3 and anti-CD15 microbeads using a QuadroMACS separator (Miltenyi Biotec, Cat. 130-090-976) and LS columns (Miltenyi Biotec, Cat. 130-042-401). Cell viability and purity of selection were confirmed by light microscopic assessment of modified Giemsa stained cytospins as detailed previously (*86*). 2 x 10^5^ cells (1 x 10^6^ cells mL^-1^) of the CD3/CD15-depleted G-CSF-mobilized PBMCs were subsequently transferred to 96U-bottom ultralow adherence plates (Nunclon Sphera, Cat. 174925) to be cultured in StemPro34 serum-free medium (Gibco, Cat. 10639011) with 1x penicillin/streptomycin (Thermo Fisher, Cat. 15140122) supplemented with stem cell factor (SCF) (R&D, Cat. 255-SC-050/CF, 20 ng mL^-1^) ± the following cytokine compositions: a) none, b) recombinant human IL-3 (R&D, Cat. 203-IL-050/CF, 25 ng mL^-1^) or c) human M-CSF recombinant protein (Invitrogen, Cat. PHC9501, 100 ng mL^-1^). A partial medium change was performed every 48 hours with 2x cytokine composites to replenish cytokines. Cell differentiation and viability were confirmed using cytospins on days 5 and 9.

### Flow cytometry analysis

Spleen leukocyte suspensions were prepared using DNAse I and collagenase D (*25*). For FACS sorting and analysis, we used previously reported protocols (*13, 19*), published HSPC definitions (*87*), indicated antibodies (see table S1), FACSCanto, LSRI, LSRII and FACSAriaIII equipment and DIVA software (BD), analyzing only populations with at least 200 events.

For human samples, an antibody panel was used to distinguish progenitors of human HSPCs (Lin^-^CD34^+^) such as common lymphoid progenitors (CLPs, Lin^-^CD34^+^CD38^-/low^CD45RA^+^), common myeloid progenitors (CMPs, Lin^-^CD34^+^CD38^+^CD45RA^-^) or granulocyte-macrophage progenitors (GMPs, Lin^-^CD34^+^CD38^+^CD45RA^+^) with mature GMPs additionally expressing HLA-DR from mature myeloid cells (either CD11b^+^CD66b^-^ or Lin^+^ ± CD14/CD16) whose IL15Rα expression was quantified. For NK cell abundance and activity, an optimized panel as published previously was used (*88*). For flow cytometry, 2 x 10^5^ cells were harvested on the day of seeding (day 0) or on days 5 and 9, respectively.

### Immunofluorescence

Freshly frozen OCT embedded (Sakura Finetek). 8 µm sections (Leica CM3050 S cryostat) were fixed 10’ in 4^°^C acetone, blocked 30’ with PBS/2%BSA, stained with 1:100 directly coupled antibody (see table S2) in PBS/2%BSA for 1 hour, mounted in ProlongGold (Invitrogen) and acquired on a LSM780 Carl Zeiss microscope.

### Microfluidic real-time RT-PCR gene expression analysis

Total mRNA extraction from 50,000 FACS-sorted cells and cDNA synthesis were performed with μMACS one step T7 template kit (Miltenyi) and specific gene expression (primers in table S3) was detected according to Fluidigm protocols as previously described (*89*) or by SybrGreen method (*13*). Ct values were calculated by BioMark Real-time PCR Analysis software (Fluidigm) using the ΔΔCt method and *HPRT* for normalization.

### Statistical analysis

Multiple statistical methods, including Student’s *t* test, Mann-Whitney *U*-test, log-rank (Mantel-Cox) test were used in this study depending on the data type, and the details can be found in the figure legends: test used and exact value of *n*. Data between two groups were analyzed with unpaired Student’s *t* tests. All statistical analyses were performed using GraphPad Prism (9.4.1). All data were expressed as medians + individual data points or means ± SEM. *P* values less than 0.05 were considered significant.

## Supplementary Materials

This file includes:

Figs. S1 to S5

Tables S1 to S3

## Acknowledgments

We thank L. Chasson and Caroline Laprie for histology, L. Razafindramanana for animal handling, M. Barad, A. Zouine and S. Bigot for cytometry support, J. Maurizio and Michaela Burkon for assistance in figure preparation. We thank Martin Bornhäuser for critical reading and discussion of the manuscript. We acknowledge the Dresden Department of Transfusion Medicine (Kristina Hölig) and the Dresden Stem Cell Laboratory (Manja Wobus and Malte von Bonin) for providing samples of G-CSF-mobilized PBMCs for the human *ex vivo* experiments. We appreciate the support of human donors of whom informed consent was obtained prior to the *in vitro* assays. We thank Philippe Pierre for *IFNAR1*-KO mice. A visual abstract, Fig. 1A, Fig. 2B, fig. S4A, and fig. S5A were created and exported with BioRender.com under a paid subscription.

## Funding

This study was supported by

Institutional grants from TU Dresden,

Institut National de la Santé et de la Recherche Médicale,

Centre National de la Recherche Scientifique and Aix-Marseille University and grants from Fondation pour la Recherche Médicale (DEQ. 20110421320) to M.H.S.,

‘Agence Nationale de la Recherche’ (ANR-11-BSV3-026-01, ANR-17-CE15-0007-01 and ANR-18-CE12-0019-03) to M.H.S.,

INCa (13-10/405/AB-LC-HS) to M.H.S.,

Fondation ARC pour la Recherche sur le Cancer (PGA1 RF20170205515) to M.H.S.,

European Research Council (ERC) under the European Union’s Horizon 2020 research and innovation program (grant agreement number 695093 MacAge) to M.H.S.

The funders of the study had no role in the study design, data collection, data analysis, data interpretation, or writing of the manuscript. M.H.S. is an Alexander von Humboldt Professor at the TU Dresden. P.K.K was partially funded by SATT Sud. J.S. is funded by the Deutsche Forschungsgemeinschaft (Clinician Scientist position, SU 1360/1-1). The authors declare no competing financial interests.

## Author contributions

P.K.K., J.S., M.D. and M.H.S. designed experiments, P.K.K. performed most experiments, C.C. contributed to initial setup and analysis of experiments. P.K.K., J.S., M.D. and M.H.S. analyzed data; C.C., J.I., and E.T. produced MCMV and helped with infection standardization; B.d.L. and M.C. contributed to processing samples for HSPC transplantation; B.d.L. tested virus and performed allogenic transplantations, J.S. and J.N performed human cell experiments; W.C, S.S., G.B., N.M.K. and C.H designed and performed gene-expression experiments; G.M., R.B. and M.F.G provided baculo-virus-produced murine M-CSF; M.D. and S.S. helped with critical reading of the manuscript; M.H.S. conceived the project, P.K.K., J.S. and M.H.S planned and managed the project, and wrote the manuscript; all authors reviewed and agreed with the final version of the manuscript.

## Competing interests

The authors declare the following potential conflict of interests: Michael Sieweke is a patent holder of WO2014167018A1 (Use of M-CSF for preventing or treating myeloid cytopenia and related complications).

## Data and materials availability

For original data, please contact.

## Supplementary materials to

**Table S1.** Information on flow cytometry antibodies. The following antibodies were used according to the manufacturer’s instructions throughout the study. Antibodies from LIVE/DEAD Fixable Aqua onwards refers to the antibodies used for the experiments using G-CSF-mobilized HSPCs. Antibodies from CX3CR1 onwards refers to the antibodies used for the allotransplantation studies.

| Antigen | Fluorophore | Clone | Manufacturer | Cat. No. |
| --- | --- | --- | --- | --- |
| CD45R/B220 | APC | RA3-6B2 | Biolegend | 103211 |
| CD11b | PerCP | M1/70 | Biolegend | 101229 |
| CD11c | A700 | HL3 | BD Pharmingen | 560583 |
| CD19 | APC | 6D5 | Biolegend | 115511 |
| CD19 | FITC | 1D3 | Biolegend | 152403 |
| CD19 | PE | 1D3 | Biolegend | 152407 |
| CD3ε | PE | 145-2c11 | Biolegend | 100307 |
| CD4 | PE | RM4-5 | Biolegend | 100511 |
| CD4 | PerCP | RM4-5 | Biolegend | 100537 |
| CD8α | Pacific Blue | 53-6.7 | Biolegend | 100728 |
| CD8α | PE | 53-6.7 | Biolegend | 100707 |
| CD8α | PerCP | 53-6.7 | Biolegend | 100731 |
| CD8β | PE | H35-17.2 | eBioscience | 12-0083-82 |
| CD49b | APC | DX5 | Biolegend | 108909 |
| Granzyme B | APC | GB11 | eBioscience | GRB05 |
| IFNβ | FITC | RMMB1 | Novus<br>Biologicals | 22400-3 |
| IFNγ | A700 | XMG1.2 | eBioscience | 56-7311-82 |
| IL-12 | APC | C15-6 | Biolegend | 505205 |
| Ki67 | Pacific Blue | B56 | Biolegend | 350513 |
| NK-1.1 | BV510 | PK136 | BD Biosciences | 563096 |
| NK-1.1 | APC | PK136 | BD Biosciences | 561117 |
| NKp46 | PE | 29A1.4 | eBioscience | 12-3351-82 |
| Streptavidin | PE-Cy7 |  | BD Biosciences | 557598 |
| Purified NK1.1 |  | PK136 | BD Biosciences | 553162 |
| Purified polyclonal rat IgG, F(ab')<sub>2</sub> fragment specific |  |  | Jackson<br>ImmunoResearch | 212-005-106 |
| CD117 | BV605 | 2B8 | Biolegend | 105847 |
| Sca-1 | PerCP-Cy5.5 | D7 | eBioscience | 45-5981-82 |
| CD34 | A700 | RAM34 | eBioscience | 56-0341-82 |
| CD16/32 | PE | 2.4G2 | BD Biosciences | 553145 |
| CD11b | PECF594 | M1/70 | BD Biosciences | 562287 |
| Ly6G | FITC | 1A8 | eBioscience | 11-9668-82 |
| Ly6C | APC | HK1.4 | eBioscience | 17-5932-82 |
| Ultra-LEAF Purified CD115 |  | AFS98 | Biolegend | 135537 |
| CD45.2 | PerCP/Cy5.5 | 104 | BD Biosciences | 552950 |
| CD45.1 | V450 | A20 | BD Biosciences | 560520 |
| Ter119 |  | TER-119 | eBioscience | MA1-70078 |
| CD71 |  | R17217 | eBioscience | 14-0711-82 |
| LIVE/DEAD<br>Fixable Violet |  |  | Invitrogen | L34955 |
| LIVE/DEAD<br>Fixable Aqua |  |  | Invitrogen | L34957 |
| CD3 | APC-Cy7 | SP34-2 | BD Biosciences | 557757 |
| CD4 | APC-H7 | SK3 | BD Biosciences | 641398 |
| CD14 | PE-Cy5 | TuK4 | Invitrogen | MHCD1406 |
| CD16 | Pacific Blue | 3G8 | BD Biosciences | 558122 |
| CD56 | PE-Cy7 | NCAM16.2 | BD Biosciences | 335809 |
| Granzyme B | PE CF594 | GB11 | BD Biosciences | 562462 |
| CD34 | PE-Cy7 | 581 | Biolegend | 343515 |
| CD38 | BV650 | HIT2 | Biolegend | 303505 |
| CD45RA | Pacific Blue | HI100 | Biolegend | 304117 |
| CD11b | FITC | M1/70 | Biolegend | 101205 |
| CD64 | PE/Dazzle594 | 10.1 | Biolegend | 305031 |
| CD66b | APC-Cy7 | G10F5 | Biolegend | 305125 |
| HLA-DR | AF647 | L243 | Biolegend | 307621 |
| IL15Rα | PE | JM7A4 | Biolegend | 330207 |
| CX3CR1 | FITC | SA011F11 | Biolegend | 149019 |
| CD45.2 | PerCP/Cy5.5 | 104 | BD Biosciences | 552950 |
| CD45.1 | V450 | A20 | BD Biosciences | 560520 |
| CD11b | BV605 | M1/70 | BD Biosciences | 563015 |
| F4/80 | BV785 | BM8 | Biolegend | 123141 |
| CD3ε | APC/AF6 | 145.2C11 | BD Biosciences | |
| Ly6C | AC7 | HK1.4 | Biolegend | 128025 |
| CD19 | PEC7 | 6D5 | Biolegend | 115519 |

**Table S2.** Information on immunofluorescence antibodies. The following antibodies were used according to the manufacturer’s instructions throughout the study.

| **Antigen** | **Fluorophore** | **Clone** | **Manufacturer** | **Cat. No.** |
| --- | --- | --- | --- | --- |
| NK-1.1<br>(IgG2a) | Unconjugated | PK136 | Invitrogen | MA1-70100 |
| m123/IE-1<br>(MCMV) | Unconjugated | IE1.01 | Capri (Center<br>for Proteomics) | HR-MCMV-12 |
| Goat anti-Mouse IgG2a<br>Cross-<br>adsorbed<br>secondary<br>antibody | Alexa Fluor 594 |  | Invitrogen | A-21135 |

**Table S3.**
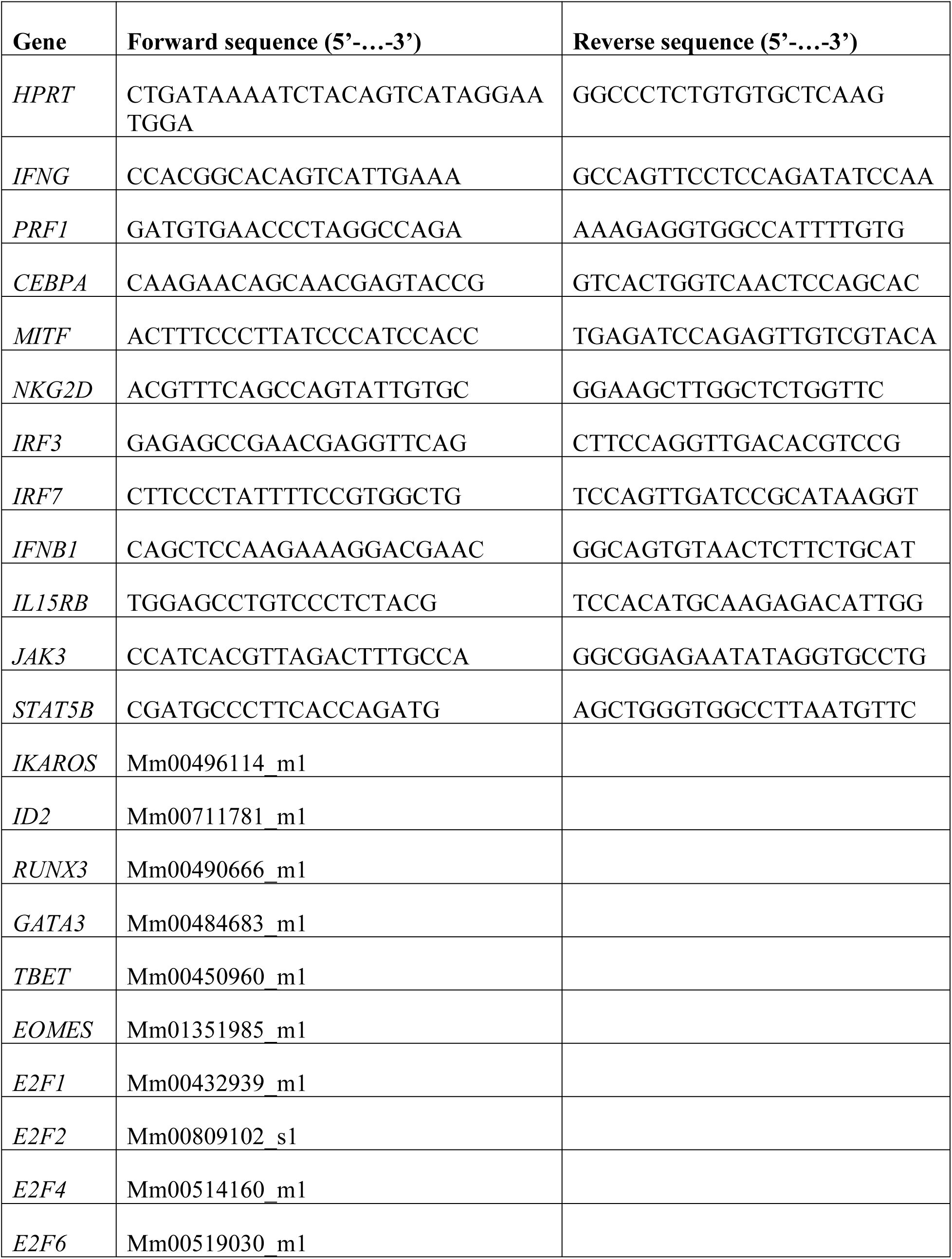
Information on primer sequences. The following forward and reverse primers were used for microfluidic real-time PCR throughout the study. The assay IDs from *IKAROS* onwards refer to Fluidigm experiments.

### Supplementary figures

**Fig. S1.**
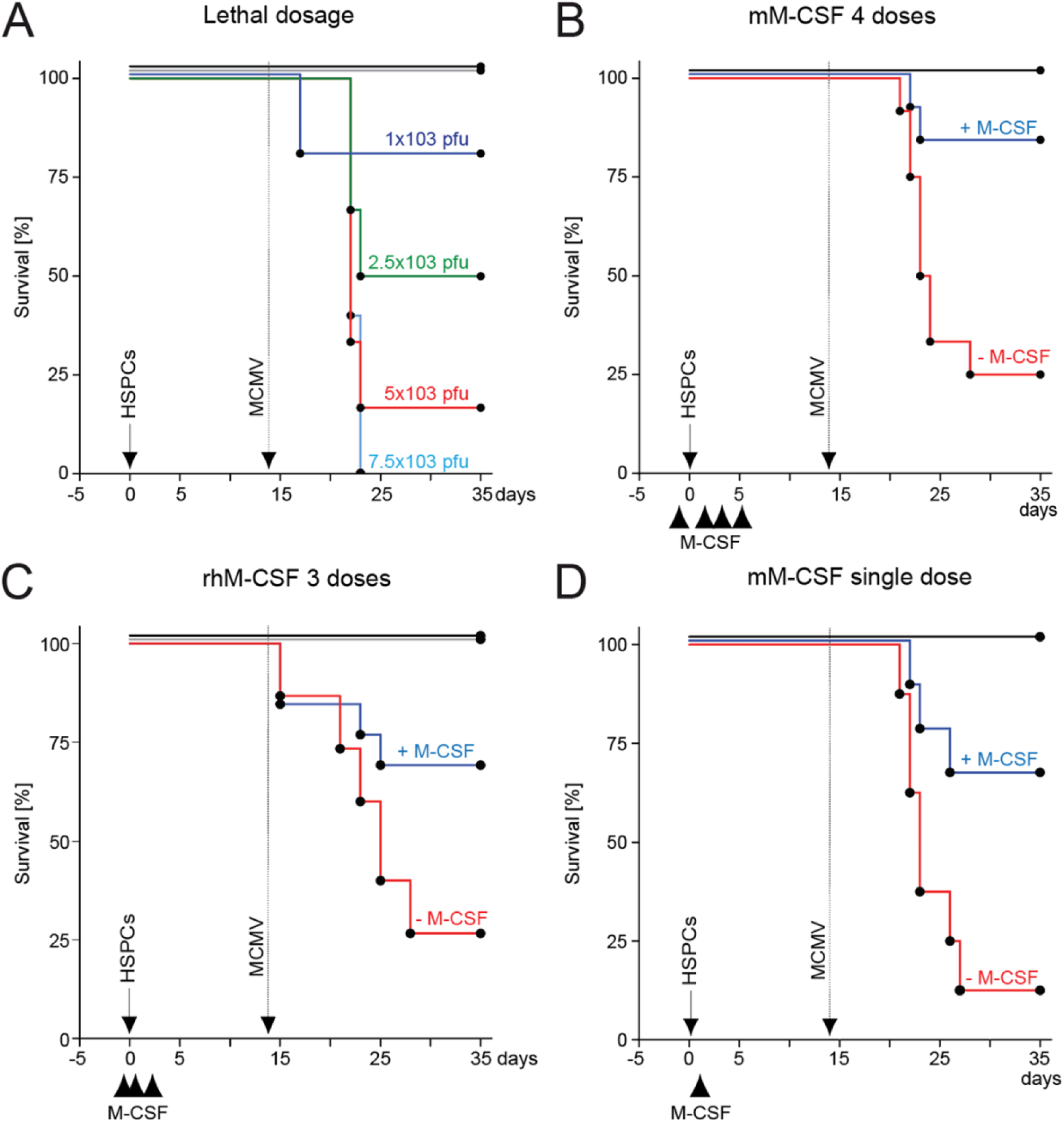
Titration of treatment regimen for MCMV infection and M-CSF treatments. (A) Survival of HSPC-transplanted mice after MCMV infection. Two weeks after HCT, mice received MCMV intraperitoneally: 1,000 PFU (violet; n = 5), 2,500 PFU (green; n = 6), 5,000 PFU (red; n = 6) and 7,500 PFU (blue; n = 5). Transplantation controls (black; n = 10). Non-irradiated, non-transplanted mice with 7,500 PFU served as infection controls (brown; n = 6). (B) Treatment regimen with different doses and sources of M-CSF. Survival of mice after infection (arrow), control (-M-CSF, red) or M-CSF (+ M-CSF, blue) or transplanted, uninfected control mice (black). HSPC-transplantation (solid arrow), MCMV infection (stippled) and different intravenous doses of control or M-CSF. Treatment regimen with 4 doses (-1h, d+1, d+3, d+5) of 10 µg baculoviral-expressed mouse M-CSF (-M-CSF, n = 12; +M-CSF, n = 11; control, n = 5). (C) Like B. Treatment regimen with 3 doses (-1h, +5h, +18h) of 10 µg human recombinant M-CSF (- M-CSF, n = 15; + M-CSF, n = 13; control, n = 5). (D) Like (B) Treatment regimen with a single dose (+5h) of 10 µg baculoviral-expressed mouse M-CSF. (- M-CSF, n = 8; +M-CSF, n = 9; control, n = 5). *** *P* < 0.0001 by Mantel-Cox test (B-D).

**Fig. S2.**
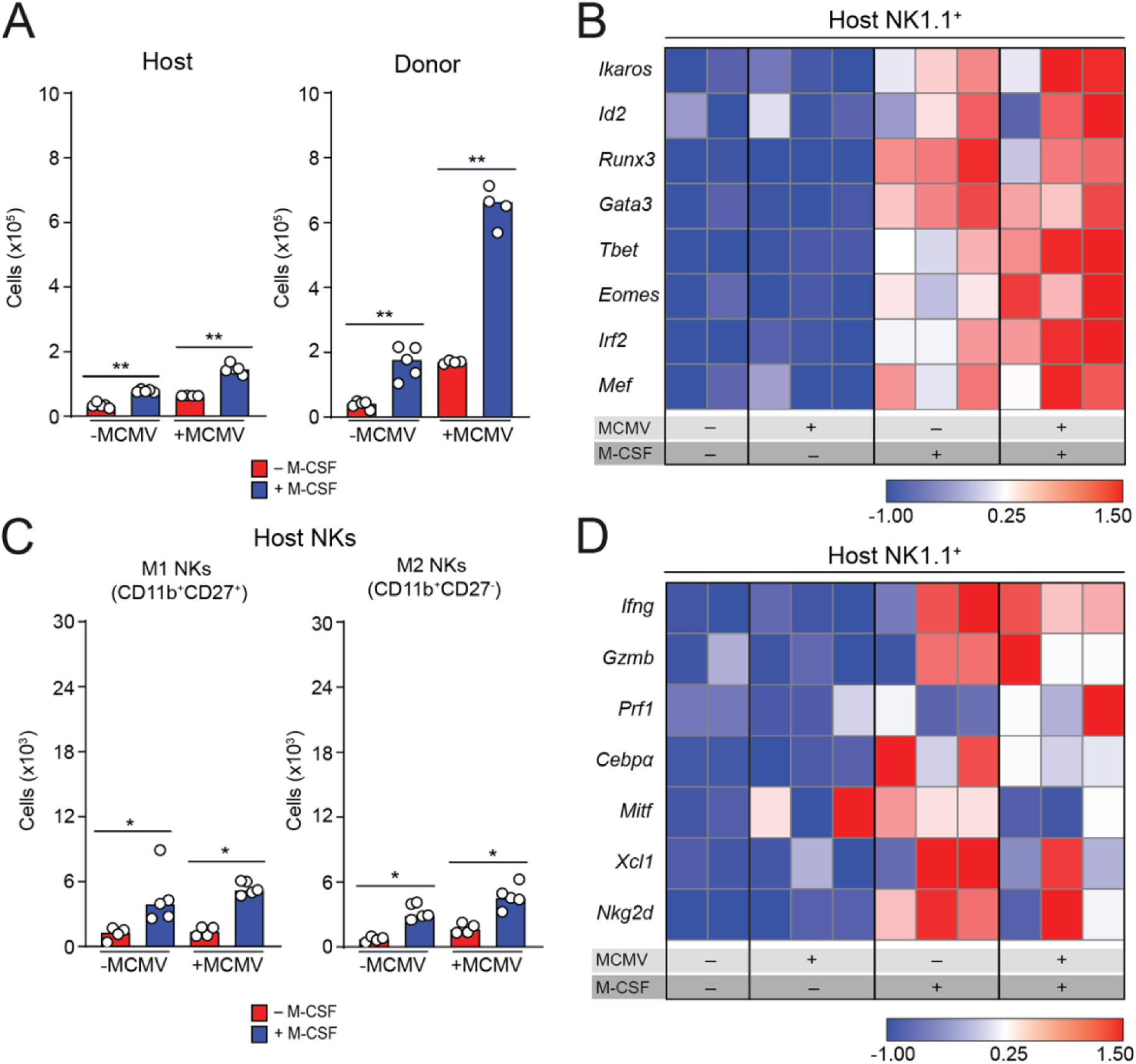
M-CSF effect on NK cell production, maturation and differentiation in donor and recipient cells after hematopoietic cell transplantation. (A) Median of absolute numbers of (CD45.2^+^) recipient (left) and (CD45.1^+^) donor (right) NK cells (CD19^-^CD3^-^Ly6G^-^NK1.1^+^). (B) Gene expression profiling of transcription factors measured by nanofluidic Fluidigm array RT-qPCR of host-derived NK cells, which were isolated from the spleens of control or M-CSF-treated recipient mice 1.5 days after MCMV of infection or time-matched, mock-infected, HSPC-transplanted mice. (C) Median of absolute numbers of host-derived M1 NK cells (CD11b^+^CD27^+^) and host-derived M2 NK cells (CD11b^+^CD27^-^) in the spleen of PBS control or M-CSF-treated recipient mice 1.5 days after MCMV or mock infection 14 days after HSPC transplantation. \**P* < 0.05 by Mann-Whitney U-test. (D) Gene expression profiling of host-derived NK cells, which were isolated from the spleen of control or M-CSF-treated recipient mice 1.5 days after MCMV or mock infection for activation and maturation related factors by RT-qPCR.

**Fig. S3.**
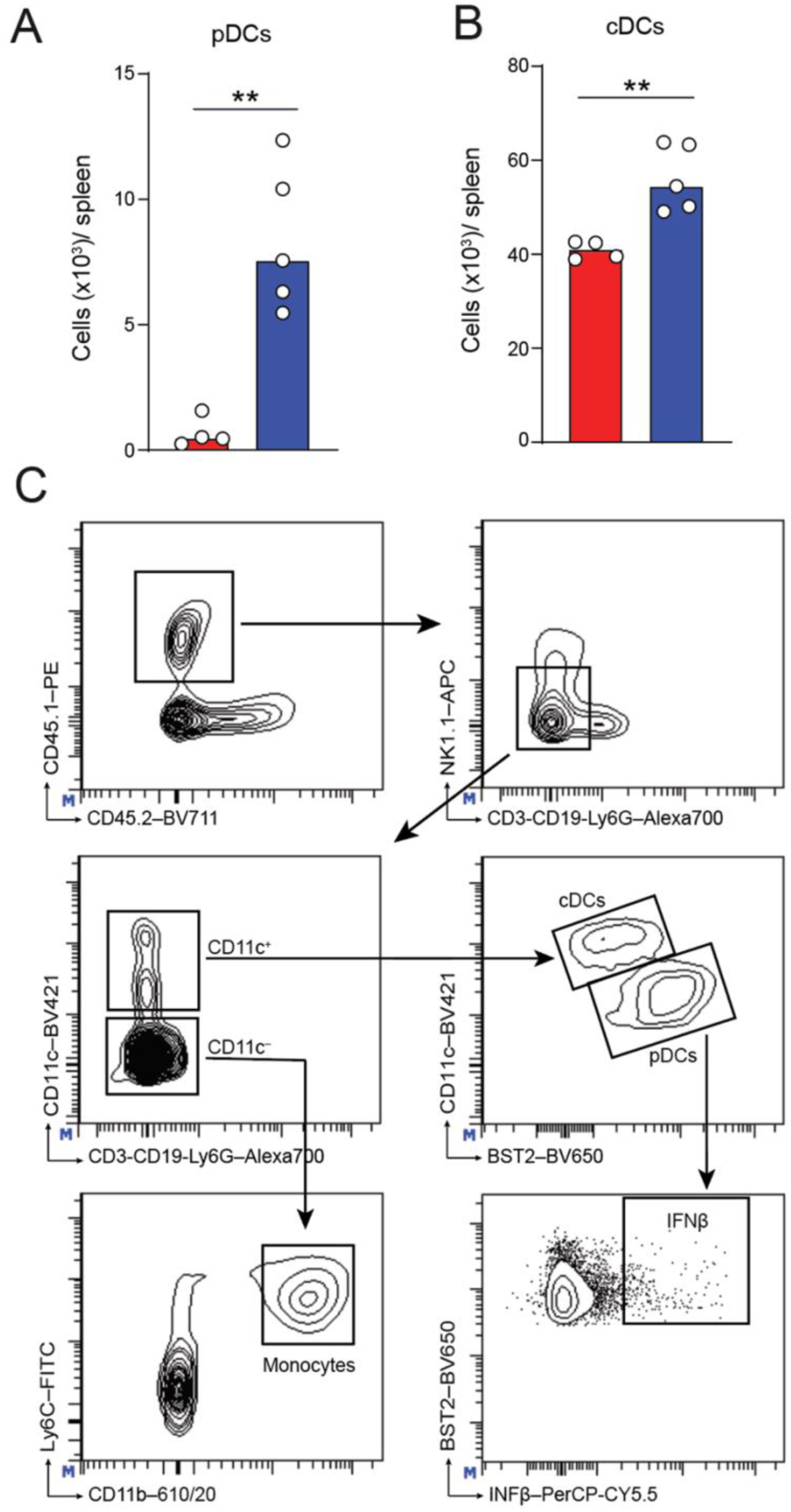
M-CSF increases myelopoiesis of plasmacytoid dendritic cells and conventional dendritic cells. (A) Median of absolute numbers of donor-derived spleen pDCs (Lin^-^ CD11c^lo^BST2^high^) of mice treated with PBS control or M-CSF 14 days after HCT and analyzed after an additional 1.5 days of MCMV or mock infection. (B) Median of absolute numbers of cDCs (Lin^-^CD11c^+^BST2^-/low^) of mice treated with PBS control or M-CSF 14 days after HCT and analyzed after an additional 1.5 days of MCMV or mock infection. (A-B) \*\**P* < 0.01 by Mann-Whitney *U*-test. (C) Gating strategy for CD45.1^+^ monocytes, pDCs, IFN-β and cDCs.

**Fig. S4.**
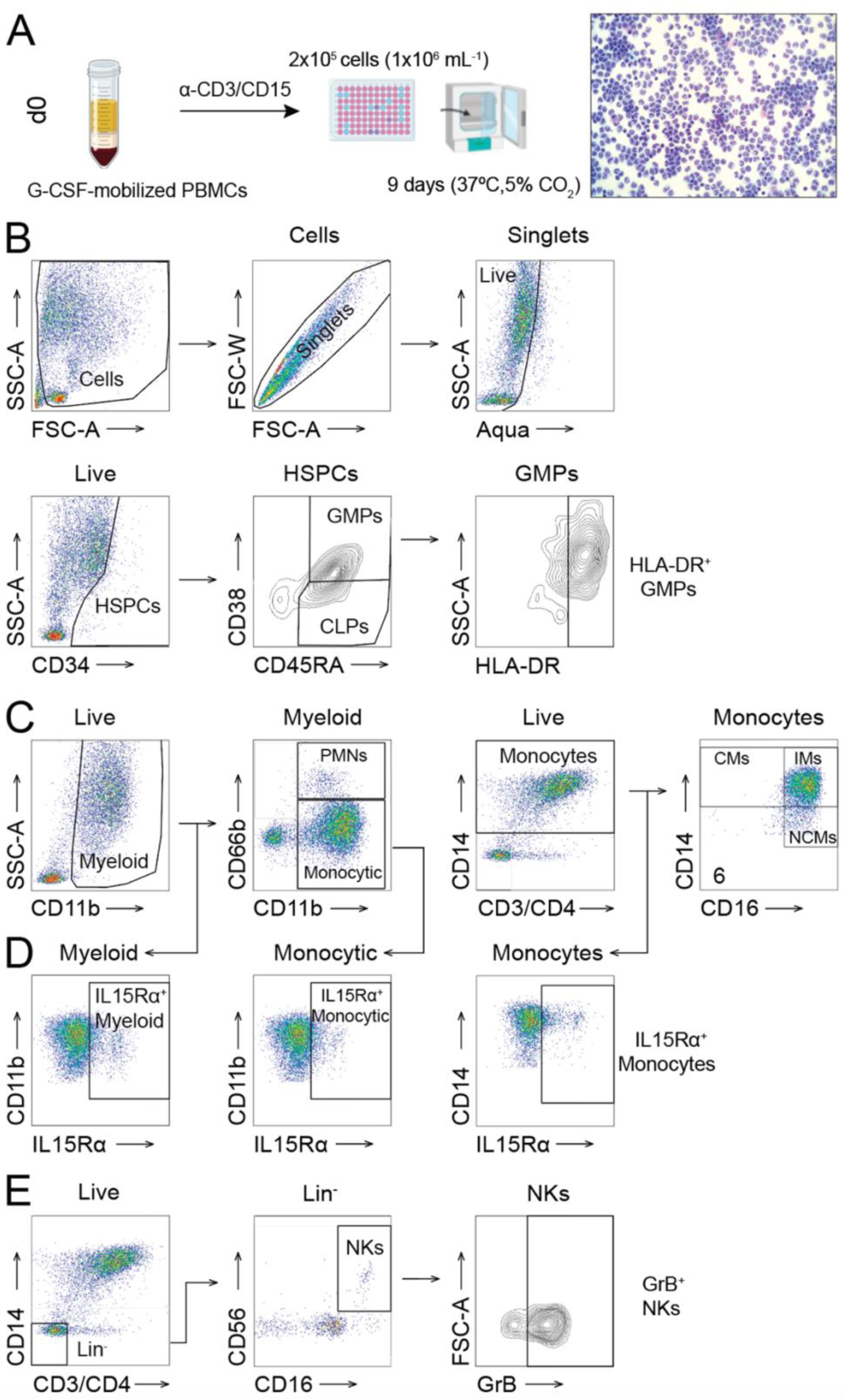
Gating strategy for M-CSF-driven myelopoiesis in G-CSF-mobilized human PBMCs, IL15Rα expression and NK cell frequency and activity. (A) Workflow for G-CSF-mobilized leukapheresis samples (B) Gating strategy for flow cytometric assessment. (C) “Live” singlets assessed for CD11b (“Myeloid” cells): polymorphonuclear neutrophils (PMNs, CD11b^+^CD66b^+^) and CD11b^+^CD66b^-^ “Monocytic” cells. “Monocytes” (CD14^+^): classical monocytes (CMs, CD14^+^CD16^-^), intermediate monocytes (IMs, CD14^+^CD16^+^) or non-classical monocytes (NCMs, CD14^-^CD16^+^). (D) IL15Rα expression measured on myeloid, monocytic cells, or monocytes identified in (C). (E) The gating strategy of OMIP-027 with minor modifications.

**Fig. S5.**
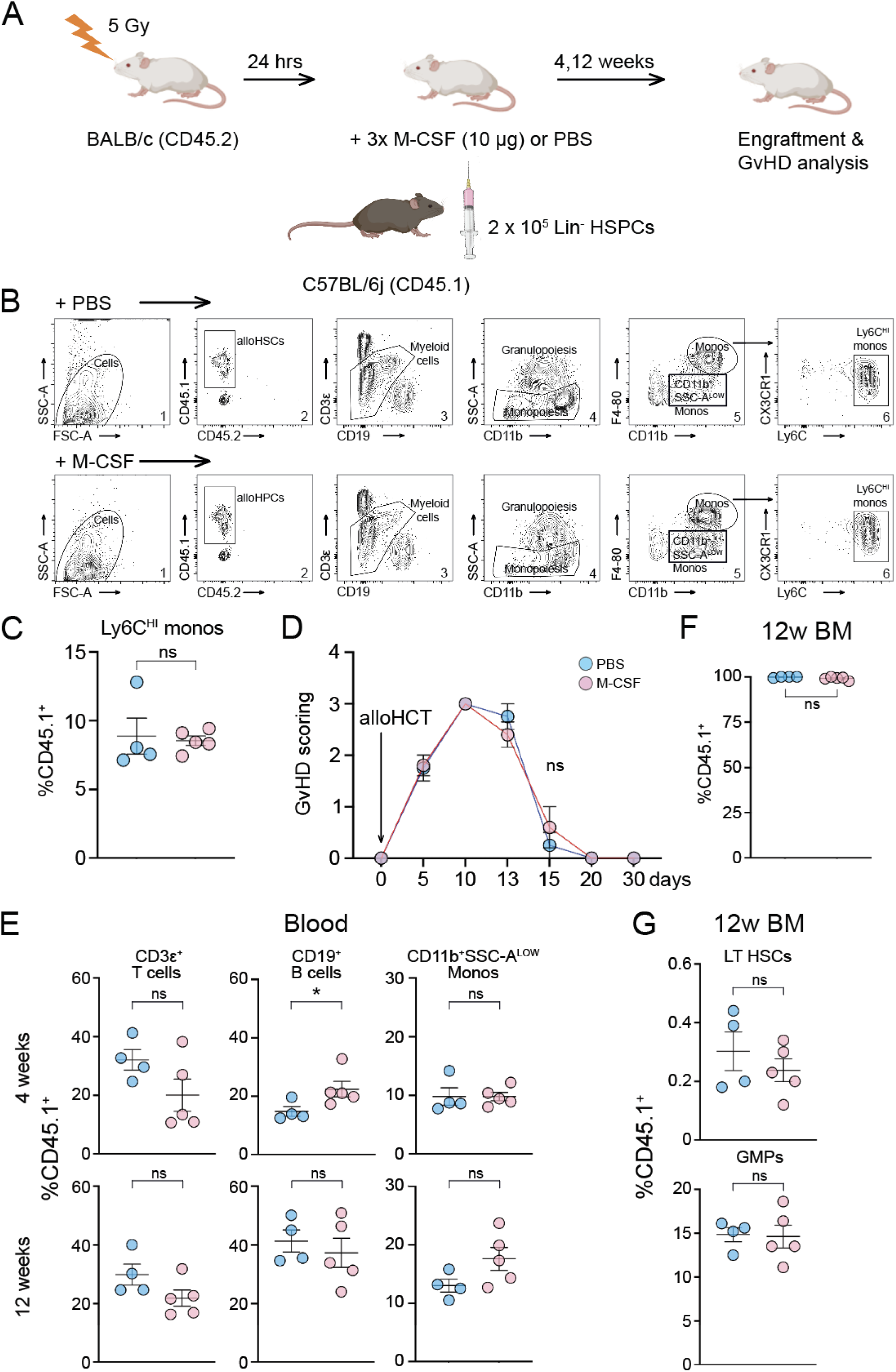
M-CSF does neither confer adverse effects on tri-lineage long-term engraftment nor on GvHD after allogeneic hematopoietic cell transplantation. (A) The protocol for allogeneic hematopoietic stem cell transplantations (alloHCT) between BALB/c CD45.2^+^ recipient and C57BL/6j CD45.1^+^ donor mice is shown. Imminently before (1 hr) or shortly after (5 hrs, 20 hrs) alloHCT with 2 x 10^5^ lineage negative (Lin^-^) hematopoietic stem and progenitor cells (HSPCs), the mice received PBS or 10 µg of baculoviral-expressed human M-CSF. Graft-versus-host-disease (GvHD) scores and engraftment of CD45.1^+^ cells were assessed at 4 and 12 weeks after alloHCT using the gating strategy by Alexander *et al*. (B). (C) Quantification of inflammatory Ly6C^HI^ CD11b^+^F4/80^+^ monocytes (monos) of CD45.1^+^ cells. (D) GvHD scoring (Lai *et al*.). (E) Tri-lineage engraftment (CD3ε^+^ T cells, CD19^+^ B cells, CD11b^+^SSC-A^LOW^ monocytes) at 4 and 12 weeks post-HCT in the blood. (F) CD45.1^+^ cells in the bone marrow (BM) 12 weeks after alloHCT. (G) Percentage of HSCs (KSL Flt3^-^CD150^+^CD48^-^) and GMPs in CD45.1^+^ lineage negative BM cells 12 weeks after alloHCT. The data are illustrated as mean ± SEM. The Mann-Whitney *U*-test was used to test for statistical significance between PBS-treated (n = 4) or M-CSF-treated allografted mice (n = 5). * *P* < 0.05, ns = not significant.

